# NIT: an open-source tool for information theoretic analysis of neural population data

**DOI:** 10.1101/2022.12.11.519966

**Authors:** Roberto Maffulli, Miguel A. Casal, Marco Celotto, Stefano Zucca, Houman Safaai, Tommaso Fellin, Stefano Panzeri

**Affiliations:** Neural Computation Laboratory, Center for Human Technologies, Istituto Italiano di Tecnologia, Genova, Italy; Department of Computer Science, Bioengineering, Robotics and Systems Engineering, University of Genova, Genova, Italy; Department of Neural Information Processing, Center for Molecular Neurobiology (ZMNH), University Medical Center Hamburg-Eppendorf (UKE), Hamburg, Germany; Department of Pharmacy and Biotechnology, University of Bologna, Bologna, Italy; Department of Life Sciences and System Biology and Neuroscience, Institute Cavalieri Ottolenghi (NICO), University of Turin, Italy; Optical Approaches to Brain Function Laboratory, Istituto Italiano di Tecnologia, Genova, Italy; Department of Neurobiology, Harvard Medical School, Boston, USA

## Abstract

Information theory provides a popular and principled framework for the analysis of neural data. It allows to uncover in an assumption-free way how neurons encode and transmit information, capturing both linear and non-linear coding mechanisms and including the information carried by interactions of any order. To facilitate its application, here we present Neuroscience Information Toolbox (NIT), a new toolbox for the accurate information theoretical analysis of neural data. NIT contains widely used tools such as limited sampling bias corrections and discretization of neural probabilities for the calculation of stimulus coding in low-dimensional representation of neural activity (e.g. Local Field Potentials or the activity of small neural population).Importantly, it adds a range of recent tools for quantifying information encoding by large populations of neurons or brain areas, for the directed transmission of information between neurons or areas, and for the calculation of Partial Information Decompositions to quantify the behavioral relevance of neural information and the synergy and redundancy among neurons and brain areas. Further, because information theoretic algorithms have been previously validated mainly with electrophysiological recordings, here we used realistic simulations and analysis of real data to study how to optimally apply information theory to the analysis of two-photon calcium imaging data, which are particularly challenging due to their lower signal-to-noise and temporal resolution. We also included algorithms (based on parametric and non-parametric copulas) to compute robustly information specifically with analog signals such as calcium traces. We provide indications on how to best process calcium imaging traces and to apply NIT depending on the type of calcium indicator, imaging frame rate and firing rate levels. In sum, NIT provides a toolbox for the comprehensive and effective information theoretic analysis of all kinds of neural data, including calcium imaging.

## Introduction

Information theory (IT), is the principled mathematical theory of communication [1]. Its use as analysis tool to measure how neurons encode and transmit information has been key to understanding brain functions such as sensation, spatial navigation, and decision-making. Mutual information (MI), the key quantity of IT, measures how well variables important for cognitive functions, such as sensory stimuli, are encoded in the activity of neurons, and how information is transmitted across brain regions. Its use has many advantages [2–8]. It provides a single-trial measure of information encoding and it is thus more relevant for single-trial behavioral or perceptual functions than trial-averaged measures of discriminability. It quantifies information in units of bits, a meaningful and interpretable uncertainty-reduction scale. It allows largely hypotheses-free measures of information encoding that place upper bounds to the performance of any decoder, and that can potentially capture the contributions of both linear and non-linear interactions between variables at all orders. Because of its generality, it can be applied to any type of brain activity recordings. Also, because neural systems may need to maximize information encoding for evolutionary reasons, applications of IT to empirical data allows a direct comparison between the predictions of normative neural theories and real neural data [5, 9]. Because of these advantages, information theory has deeply influence neuroscience over many years [5, 7, 9–13].

Earlier work using information theory to analyze empirical neuroscience data has focused on low-dimensional measures of neural activity such as such as single neurons, small neural populations or aggregate measures such as LFPs/EEGs (because of the systematic errors in estimating information with the small numbers of trials that can be collected empirically are exacerbated with high-dimensional neural responses [14]). It has also focused mostly on information encoding, regardless of the downstream use of the encoded information. Seminal studies of this kind have used electrophysiological recordings of neural activity to demonstrate the role of single-neuron spike timing for the encoding of sensory information [6, 7, 15–17]. Other studies have provided the foundations of how trial-to-trial correlations between neurons shape the encoding of information and create redundancy and synergy in pairs of neurons [18–20]. Further studies have examined how information is encoded in the neural oscillations found in aggregate measures of neural activity such as Local Field Potentials (LFPs) [21, 22]. Several algorithms have been proposed for the application of IT to these low-dimensional neural data [6, 23, 24]. Their ability to provide accurate and data-robust information estimates has been extensively validated and demonstrated on electrophysiological recordings, including on spike trains of small populations and on LFPs and EEGs [24–27], and their use and dissemination has been aided by software toolboxes [25, 28–32].

Over the last decade, due to major progress in the simultaneous recording from many neurons and/or brain areas, and in the measure and quantification of behavior [33], neuroscience research [34–36] – and consequently neuroscientific IT – has evolved to investigate how behavior and information processing emerge from the interaction and communication between neurons and across brain areas. For example, recent work has coupled IT with dimensionality-reduction techniques to study how information is encoded in populations of tens to hundreds of neurons [37–46], and of how patterns of synergy between pairs of neurons are organized within larger networks [20]. Studies have also characterized the transfer of information between neural populations [47, 48] and between brain areas [49–51]. Importantly, neuroscientific IT has also been used to measure the information carried by neural activity not only about sensory stimuli, as in traditional studies, but also about behaviorally relevant signals such as choice and reward [45]. Moreover, Partial Information Decompositions (PID) [52] has extended Shannon’s IT to quantify how much of the information encoded in neural activity is used to inform behavioral choices during perceptual discriminations [53, 54] and synergistic or redundant transfer of information across brain regions [49]. However, progress in using latest IT advances in neuroscience to address large populations, behavioral relevance and information transmission, synergy and redundancy with PID, has been slowed by the absence of comprehensive toolboxes including all or most these recent tools.

Key to the recent progress in understanding the relevance of neural population activity for behaviors has been the application of 2-Photon (2P) fluorescence microscopy [55–57] to image the activity of populations of neurons in animals performing cognitive tasks [58–63], even over days or months [64–68]. However, applying information theory to 2P imaging recordings is particularly challenging. 2P calcium imaging measures neural activity only indirectly (by the optically recorded fluorescence signal changes that originate from changes in calcium concentrations related to changes in neural activity), and it generally has low SNR and limited temporal resolution. Understanding how to optimize the use of information theory to analyze large-scale recordings of populations with 2P imaging during behavior would greatly aid progress in studying neural population coding.

Here, we introduce the Neuroscience Information Toolbox (NIT) to specifically address both the need of having a single open-source toolbox including many recent advances in IT tools for neuroscience and pf optimizing its use for 2P calcium imaging. NIT provides a comprehensive set of IT tools (including MI, directed communication measures, PID tools, binned and copula probability estimators, and limited sampling bias corrections) applicable to both discrete and continuous measures of neural activity. It thus can be used with both direct electrophysiological recordings of action potentials and with indirect measures of neural activity, such as LFP, EEG, fMRI and 2P imaging. Algorithms that we implemented and optimized in NIT were already validated on electrophysiological recordings [25–27]. However, here we study extensively, both with realistic simulations and with analysis of real data, how best to extract from 2P imaging data information about variables of interest (sensory stimuli, behavioral choice, and/or the underling firing levels of neurons) and how best to tune algorithms for information measures and for calcium imaging processing depending on factors including imaging frames, calcium indicators, signal-to-noise ratio of fluorescence and neural firing regimes.

## Results

### NIT: a complete toolbox for information theoretical analysis of neural data

We present NIT, the Neuroscience Information Toolbox. NIT is a comprehensive package of open-source tools for information-theoretical analysis of neuroscience data. NIT is fully documented, and its MATLAB interface allows easy integration with custom built analysis pipelines.

Features and structure of NIT are shown in **Figure 1**. At the core of the software sits a set of modules for the calculation of information theoretic quantities. The software consists also of a set of routines for applying dimensionality reduction and neural decoding strategies. Some of the computations are performed through *ad-hoc* developed interfaces to external libraries which are distributed with the code, making NIT a self-contained toolbox. The key features and functions of the software are briefly described in the following sections.

**Figure 1.**
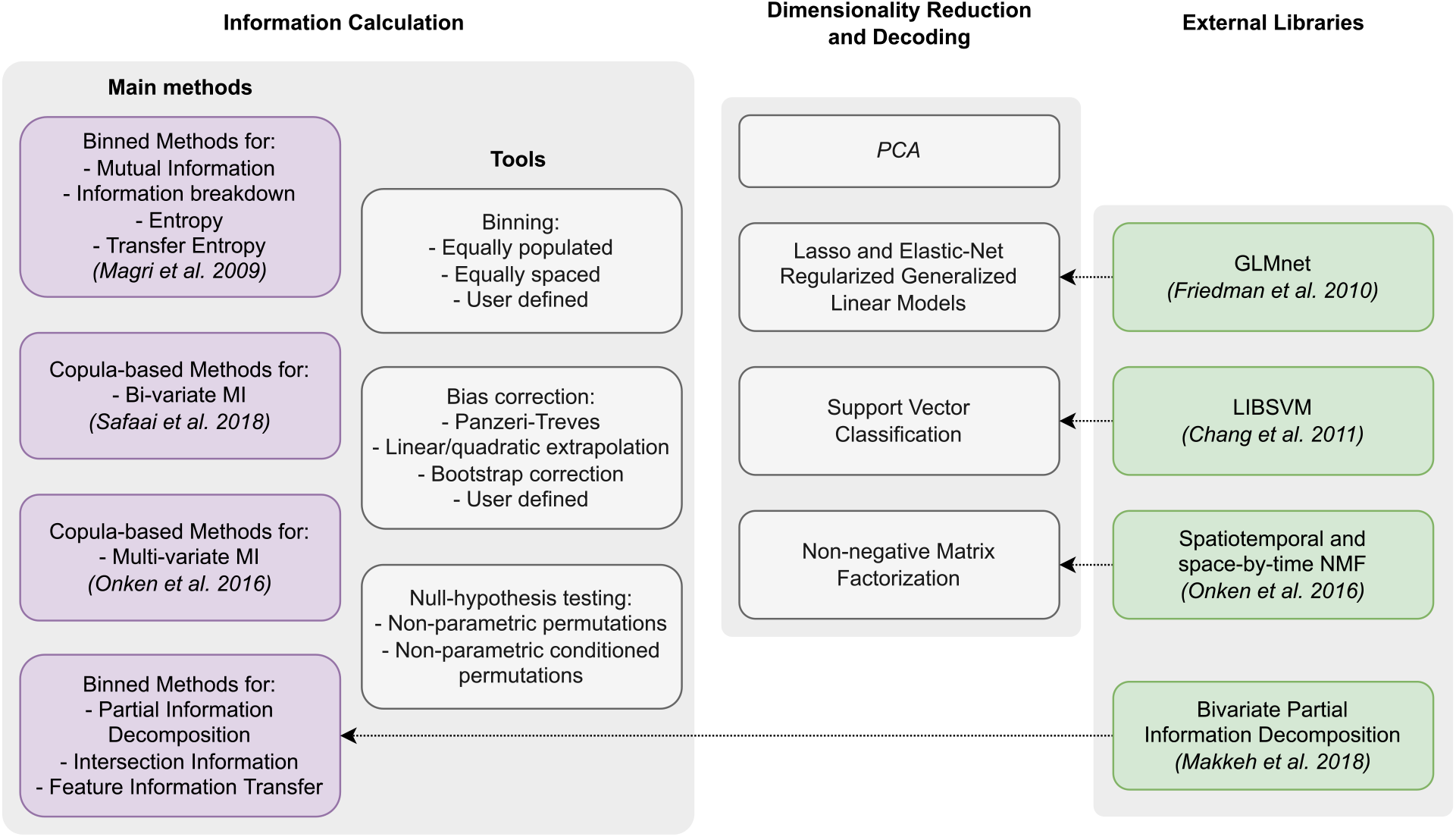
Structure of Neuroscience Information Toolbox (NIT) The toolbox comprises modules (black boxes) for calculation of information-theoretic quantities and dimensionality reduction. External libraries (green boxes) are interfaced (arrows) with some of NIT native modules to integrate their functionalities.

In the following, we first list and explain the various information theoretic functions and features included in the toolbox. We then introduce the detailed simulations of 2P calcium imaging recordings together with the results of the parametric study used to discuss the limitations of extracting information from those data as opposed to electrophysiology. Finally, we apply NIT to experimental data, first to validate what we have observed on synthetic data, as well as to illustrate how the methods implemented in NIT can be effectively used to reveal a higher level of detail of the information processing principles in the brain.

### Information theoretic algorithms and functions implemented in NIT

#### Mutual Information

MI between two random variables *R* (in this example the neuronal response) and *S* (in this example an external stimulus) measures how well a single-trial knowledge of one variable reduces our uncertainty about the value of the other variable is defined as follows [1]:

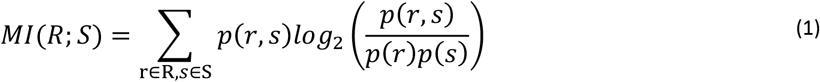

where *p*(*s, r*) is the joint probability of observing in a given trial stimulus *s* and response *r*, and *p*(*s*), *p*(*r*) are the corresponding marginal probabilities. *MI*(*S*; *R*) is measured in units of bits, it is nonnegative and it is zero if and only if *S* and *R* are statistically independent. One bit of information means that the knowledge of one variable halved the uncertainty about the other variable., *R* can be either univariate (e.g. time-averaged single neuron activity) or multivariate (e.g. neural population activity, with each dimension of *R* quantifying the activity of each neuron in a population). NIT accepts either univariate or multivariate entries for both responses and stimuli (useful when several stimulus features are varied across trials). The value of MI is computed once these probabilities are measured from the data over repeated experimental trials and inserted into Equation (1). Different methods to compute MI from real data typically differ depending on how these probabilities are estimated from the data. Three different MI calculation methods are provided in NIT.

The first one, the direct or plug-in method, consists in estimating the probabilities in Equation (1) by simply counting the number of occurrences of the discrete values of both *R* and *S* across repeated presentations of the stimulus. The plug-in method does not make assumptions on the shape of the probability distributions and has a low computational cost. To make the plug-in method applicable to cases in which *R* and/or *S* are continuous (e.g. *R* will be continuous if is extracted from unprocessed 2P calcium traces or from LFP traces), NIT has two built-in discretization functions, that bin data in equally-populated or equally-spaced classes. Equally-populated binning maximizes the entropy available in the neural response for a given number of bins and thus often leads to larger information values, whereas equally-spaced binning preserves the shape of the original probability distribution. An interface is provided for inserting into the workflow other user-defined binning methods.

A second method, applicable only when the underlying distributions of the data are Gaussian, relies on fitting a Gaussian probability density function to the data. This method, suitable for continuous data not discretized in post-processing, is less prone to limited sampling bias (see below) than the direct plug-in method. However, it is applicable only when signals are approximately Gaussian. This may hold in specific instances for aggregated electrical signals (LFP, EEG, MEG) [21, 25, 30], but it does not hold for 2P calcium traces of individual cells [69].

Finally, NIT implements also a Copula estimator, including both parametric Copulas [30, 70] and Non-Parametric Copula (NPC) MI estimation [71]. Joint multi-dimensional probabilities distributions can be expressed in terms of marginal probabilities and a copula, a mathematical term that specifically describes the statistical dependences between the variables (see Materials and methods). The MI between two variables depends on the copula but not on the marginal probabilities. This allows to estimate MI without calculating the latter [30, 70, 71]. In the NPC approach, copulas are estimated non-parametrically with Kernel methods rather than with parametric forms, allowing largely assumption-free information estimations and avoiding potential mis-estimations of information due to wrong parametric assumptions being used [71]. Estimating MI with NPC has a much higher computational cost compared to the direct plug-in method, at the advantage of being more accurate and not requiring the discretization of continuous variables (although it can be applied also to discrete variables). As an alternative, we also implemented parametric copula estimator, which use parametric assumptions for the joint probability density estimators. This has an advantage in terms of computational costs but it may become highly inaccurate when the Gaussian assumptions are not met [71]. For continuous margins, we provide implementations of the normal and the gamma distributions. For discrete margins, we provide the Poisson, binomial and negative binomial distributions. As bivariate copula building blocks, we provide the Gaussian, student and Clayton families as well as rotation transformed Clayton families [70].

#### Mutual Information breakdown to quantify the information content of neuronal correlations

The information about the stimulus encoded in the activity of a population of individual neurons depends on the strength and structure of correlations among neurons [8, 35]. NIT allows to quantify how correlations affect neural population encoding of the stimulus by using the Information Breakdown formalism [19]. The MI between the stimulus and the neuronal population response *R* (a multi-dimensional vector containing the activity of each neuron in a given trial) is divided in components that capture the different ways in which correlations affect neural population information, as follows:

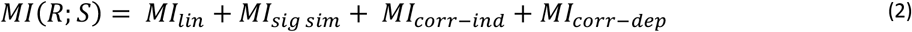

where *MI_lin_*, the linear term, is simply the sum of the *MI* about the stimulus carried by the individual neurons. The other terms, capturing the differences between *MI*(*R*; *S*) and *MI_lin_* reflect the effect of the statistical dependencies between neuronal responses. Such dependencies are traditionally conceptualized as signal correlations (correlations of the trial-averaged neural responses across different stimuli, quantifying the similarity of tuning to stimuli of different neurons) and noise correlations (correlations in trial-to-trial variability of the activity of different over repeated presentations of the same stimulus, quantifying functional interactions between neurons after discounting the effect of similarities in stimulus tuning), see e.g. [35, 72, 73]. The term *MI_sig sim_*, always less than or equal to zero, quantifies the reduction of information (or increase in redundancy) due to signal correlations (that is, because neurons have partly similar response profiles to the stimuli). *MI_corr–ind_*, a term that can be either positive or negative, quantifies the increment or decrement of information due to the relationship between signal correlation and noise correlation. The term is positive (providing synergy) if signal and noise correlations have opposite sign, while is negative (providing redundancy) if signal and noise correlations have the same sign [19]. *MI_corr–dep_* is a non-negative term that quantifies the information added by the stimulus modulations of noise correlations [19]. The information breakdown includes as a sub-case other types of decomposition and quantifications of the effect of correlations in population activity. For example, *MI_corr–ind_* + *MI_corr–dep_* quantifies the total effect of noise correlations on stimulus information and equals the quantity Δ*I_noise_* defined in [74]. Similarly, *MI_lin_* + *MI_sig–sim_* quantifies the information that the population would have if all single neurons properties were the same but noise correlations were absent, and equals the quantity *I_no–noise_* of [74]. Finally, *MI_corr–dep_* equals the quantity Δ*I* introduced in [75] as an upper bound to the information that would be lost if a downstream decoder of neural population activity would ignore noise correlations. The information breakdown formalism and the related quantities that can be obtained from it have been used in many studies to empirically characterize the effect of correlations [8, 16, 20, 38, 76–79].

#### Partial Information Decomposition

Other methods to decompose the contributions of multivariate dependencies between neurons to information carried by populations include the Partial Information Decomposition (PID) [52]. In the form implemented in NIT, PID is applied to three stochastic variables (*R*_1_, *R*_2_, *S*) (e.g. two neurons with responses *R*_1_ and *R*_2_ respectively, and a stimulus variable *S*). The method decomposes the information that two of them (called source variables, in the example above the two neuronal responses) carry about the third one (called target variable, in the example above the stimulus), in four non-negative and well-interpretable terms called “atoms”, as follows:

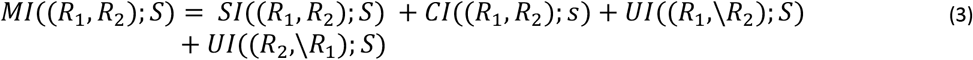

In Equation (3): *SI*((*R*_1_, *R*_2_);*S*) is the shared (redundant) information that *R*_1_ and *R*_2_ carry about *S*; *UI*((*R*_1_,\*R*_2_); *S*) is the unique information about *S* that is carried by *R*_1_ but is not carried by *R*_2_; *UI*((*R*_2_,\*R*_1_); *S*) is the unique information about *S* only present in *R*_2_ but not in *R*_1_; and *CI*((*R*_1_, *R*_2_);*S*) is the complementary (synergistic) information about *S* that is available only when *R*_1_ and *R*_2_ are measured simultaneously. NIT calculates the above PID three-variate decomposition using the so-called BROJA definition [80] through a specifically designed interface to the BROJA-2PID algorithm [81].

#### Intersection Information

One application of PID is the measure of Intersection Information (*II*, see [82, 83]). II applies to tasks such as perceptual decisions in which in each trial a stimulus (*S*) is presented, neural activity (*R*) is recorded and the subject’s perceptual report of which stimulus was presented is measured as a behavioral choice (*C*). *II* measures, in bits, how much of the stimulus information carried by neural activity *MI*(*R*;*S*) is used to inform the behavioral choice, and is defined in terms of PID as follows [83]:

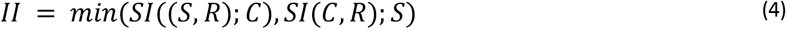

As shown in Ref [83], this expression quantifies the part of information carried by neural activity that is shared between stimulus and choice, and that at the same time is part of the overall information between stimulus and choice. *II* is non-negative, is bounded by the stimulus and choice information carried by neural activity, and by the information between stimulus and choice. *II* has been used in several studied to determine the behavioral relevance of aspects of neural population codes (e.g. [39, 54, 83]). NIT has a specifically built module for the calculation of *II* with the plug-in probability estimation method.

#### Measures of directed information transfer between neurons or brain regions

NIT implements also the most used information-theoretic measure of directed information transfer between different brain regions or neurons: Transfer Entropy (TE) [84], equivalent under the definition we use to Directed Information [85]. TE is an information-theoretic measure of the causal dependency between the time series of a putative sender *X* and the time series of a putative receiver *Y*. It is based on the Wiener-Granger causality principle, stating that a signal *Y* is causing *X* if the knowledge of the past of *Y* reduces the uncertainty about the future of *X*. Given the time series *X* and *Y* of two signals simultaneously recoded over time from different neurons or brain regions, TE is defined as:

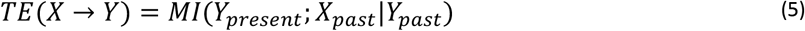

Where *Y_present_* is the value of signal *Y* at the present time, and *X_past_* and *Y_past_* are the values of *X* and *Y* at a set of *k* past times. TE computes the MI information that the past values of *X* carries about the present value of *Y*, discounting the information that the past of *Y* carries about its own present value. These measures of directed information transfer have been widely used to characterize communication between brain regions (see e.g.[47, 48, 50, 86]).

NIT allows calculating TE using the direct plug-in method. It allows to define the set of k past value used to compute TE. In most applications, TE is computed using one past value for X and Y, defined by the delay between the selected past value and the present [48, 87, 88]. However, NIT allows to include past values over a range of different delays from the present. NIT features also an optimized routine for fast calculation of TE on spike trains, taking advantage of the reduced probability space deriving from binary signals [89].

Note that NIT implements also other more recent extensions of directed information calculations derived from the PID. For example, it implements also the recently introduced Feature-specific Information Transfer (FIT) [90]. FIT extends the previously described TE by computing not only the total amount of directed information that is transmitted from the putative sender *X* and receiver *Y*, but quantifying how much of this total transmitted information relates to a specific stimulus feature of interest *S*.

Conceptually, FIT quantifies how much of the MI encoded by the present activity of *Y* was shared (redundant) with information about *S* present already in the past of *Y* while being unique with respect to the stimulus information that was encoded by past activity of *Y* [90].

Importantly, NIT allows computing also other more refined directed information transfer measures derived from PID which can be expressed in terms of appropriate combinations of MI quantities, such as those introduced in Refs [49, 91].

#### Limited sampling bias corrections

Accurate estimation of information quantities depends on accurate estimation of probabilities. Measuring probabilities from a limited number of experimental trials leads to statistical fluctuations in the estimated probabilities, which in turn leads to both statistical and systematic errors in information measures. The systematic error, or limited sampling bias, is due to the non-linear dependence of the information on the probabilities [14]. In most conditions, the limited sampling bias is positive, meaning that limited sampling tends to overestimate the MI [14, 92]. Intuitively, this is because differences of stimulus-specific neural response probabilities generated by random fluctuations due to limited sampling result through the MI equation as genuine, information -bearing features. The amount of bias is typically higher for less informative variables, and it decreases approximately linearly with the number of trials [14, 93]. Thus, although the limited sampling bias is present in all calculations of MI, it is particularly prominent for neuroscience experiments because of the limited number of trials that can be collected and because of the relatively small information values of neural activity (in our experience, in typical experiments with subjects performing tasks while recording brain activity, it is extremely rare than more than ~100-20 trials per stimulus or task condition are available, and information values of individual neurons are usually much smaller than one bit).

Fortunately, several bias correction procedures have been developed, with reduce substantially the limited sampling bias from neural measures. In case of stimulus-response information *MI*(*S*; *R*), Equation (1), most measures work well when the number of trials per stimulus is at least 4-10 larger than the number of possible values of response *R* [14, 23, 28]. This is a rule of thumb that is useful to set the number of bins used to discretize the neural response R. NIT is equipped with a sets of well-used for limited sampling bis correction in MI measure: Panzeri-Treves [23], linear and quadratic extrapolation [94], the shuffling procedure [14], the Best Upper Bounds (BUB) estimator[95], and the bootstrap correction [96]. An analytical bias correction method is specifically available for the Gaussian method [25]. Interfaces for easy plug-in of user-defined bias correction routines are available. A complete list of the compatibility between information-theoretic measures, bias correction strategies and information estimation methods implemented in NIT is provided in **Supplementary table 4**.

One point of interest that we found while running the NIT on simulated data is that, while the size of the limited sampling bias for mutual information follows well the analytical predictions of analytical polynomial expansions of the bias in terms of the inverse of the numbers of trial (e.g. [14]), the bias of II (which is not a mutual information quantity, but only a part of a mutual information quantity) was in general smaller than that predicted for mutual information with the same numbers of trials and response binning. In measures comparing mutual information with PID or II quantities, we thus recommend (as we did in **Figure 8**) to evaluate and compare the bias of PID and mutual information quantities in stretches of data in which we know information must be null (e.g. pre-stimulus time windows for stimulus information or II) and use those as estimates of bias values.

When analyzing multi-dimensional data (e.g. the simultaneous responses of neurons in a population), the number of possible responses of the population increases exponentially with the size of the population. For example, the binary activity of a population of 10 neurons recorded simultaneously can take 2^10 states, which would require an unrealistic number (~10000) of trials for accurate limited sampling correction. This makes it impossible to compute directly information from large populations [14, 97]. Dimensionality reduction and neural decoding algorithms, several of which are embedded as modules in NIT (**Figure 1**) embedded in NIT allow to analyze highly multi-dimensional data with a limited amount of trials.

#### Dimensionality reduction and neural decoding

Dimensionality Reduction (DR) methods are a precious tool for performing information-theoretical analyses of multi-dimensional neural data, as they allow to reduce the dimensionality of the response space *R* in a meaningful way at the expenses of small information losses.

Within NIT we implemented, and coupled with the information theoretic calculation, many such DR methods that have been popular in the analysis of neural activity. The pipeline first maps the multi-variate neuronal response *R* to a lower-dimensional space 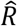, then NIT computes the mutual information 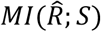 between the reduced neural variables 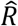 and *S*. The compression of the neural response space cannot increase the information and may lead to some information loss because of the data processing inequality [98]. However, it allows a more reliable sampling of the probability space with the limited number of experimental trial available.

The first class of DR methods implemented in NIT can be described as supervised decoding methods. These methods predict in each trial the most likely value of the stimulus S that was presented given the observation of the neural response *R* in that trial. This data compression for information calculations is popular [38, 39, 44] as effectively it reduces the response *R* to the smallest space that can in principle preserve all information about *S* (that is, the *S* space itself). Two modules for neural decoding, implementing high popular decoding methods in neuroscience, are provided in NIT. The first one is based on linear, logistic or multinomial regression through elastic-net penalized Generalized Linear Models (GLM). The core of the GLM regression functionalities are provided by the GLMnet [99] library, directly interfaced with NIT. This ensures fast and reliable decoding on large datasets characterized by sparse neuronal activity. Such types of decoders have been popular for neural activity analysis [39, 100, 101]. A second method for neural decoding applies a Support Vector Machines (SVM) for multi-class classification, which is also popular in neuroscience [102–104]. The back-end for SVM classification in NIT relies on the LIBSVM [105] package, providing fast implementation for multi-class Support Vector Classification and Regression.

NIT contains two modules for applying dimensionality reduction strategies that compress the space of neural responses in an unsupervised way without relation to the structure of the stimulus. The first one performs Principal Component Analysis (PCA), often used in neuroscience [106], through a custom-built fast MATLAB implementation. A second method is based on a Space-Time Non-negative Matrix Factorization (STNMF) [107]. The method, specifically designed for the analysis of spike trains, allows to decompose the neuronal response through a space-by-time tensor factorization. Moreover, it identifies ensembles of simultaneously active neurons and the temporal profiles of their activity. STNMF has been successfully used to extract information-rich features from the neural activity [107].

#### Hypothesis testing

NIT also provides algorithms to test the hypothesis that the measured information values are significantly different from a null hypothesis distribution of null information. While plug-in values of information for asymptotically large number of trials follow a chi-square distribution and their significance could be tested parametrically, no parametric null hypothesis distribution is known for finite number of trials (as it always the case in real calculation) and for methods different from plug-in. The well-established method to test for the significance of mutual information is the non-parametric permutation test in which all or part of the data structure is randomized to remove its information content [25, 30, 38, 108, 109]. This test computes, from many different random permutations of the data, a null-hypothesis distribution and a significance threshold to test that hypothesis that a measured value of information (which could be non-zero because of sampling bias or statistical fluctuations even if the data contain no information) for significance of information given the number of trials available and computational method used.

Significance for the value of *MI*(*S;R*) is computed by randomly permuting (or “shuffling”) the neural response *R* across experimental trials to destroy all information they carry about *S*. When computing multivariate information measures, it is sometimes of interest to test the significance of values of information between two variables conditioned on the value of other variables. For example, whether the activities of two neurons *R*_1_ and *R*_2_ have statistical dependences beyond the one induced by the common tuning to the stimulus S, can be tested by computing the significance of *MI*(*R*_1_; *R*_2_\*S*), the conditional mutual information between *R*_1_ and *R*_2_ given S. Whether R2 carries stimulus information not carried already by R1 can be tested by computing the significance of *MI*(*R*_2_;*S*\*R*_1_), the stimulus information of R2 conditioned on R1 [110]. Significance testing of information values conditioned or partialized on values of other variables can be more precisely done by shuffling the statistical relationship between the variables we compute information about at fixed value of the variables we condition upon [110, 111]. This conditioned shuffling destroys the relations between the variables we compute information about while preserving the relationship that each of them individually has with the variables we condition upon.

In NIT, we implemented routines that easily create null-hypothesis distributions and significance thresholds for both standard and conditioned mutual information values, performing shuffling of any variable possibly at fixed values of other variables, with the number of different shuffles created a parameter of the analysis.

### Extensive validation of NIT on simulated 2P data

NIT is a general-purpose toolbox, usable on any kind of neuroscientific data. The above-described algorithms implemented for computing information from neural activity have been extensively used and highly validated over the years with electrophysiological recording of spiking activity of single neurons and populations and with aggregate electrical measures of neural activity such as LFPs and EEG [14, 25, 26, 112–115]. As a result, we known well how to set the parameters of information theoretic calculations with such signals. However, studies of how best to apply these methods to 2P calcium imaging data are still limited, and no systematic validation is available.

Thus, we next validated the capabilities of NIT to extract stimulus information from 2P calcium imaging experiments through extensive simulations of synthetic 2P traces. In the analysis, we strived to cover a wide range of experimental conditions, relating both to the neuronal response and its modulation by the stimulus as well as the experimental apparatus. We first detail the model for the generation of imaging traces, followed by testing the algorithms in NIT in an extensive parametric sweep across all conditions examined. Aim of this effort was to offer a solid validation on how to analyze 2P data using information theory, highlighting the difference between the information content in imaging data compared to traditional electrophysiology analysis, as well as the advantages of non-parametric copula over binned estimators when applied to imaging data.

#### Forward model for the generation of synthetic fluorescence traces

To quantify the extent to which we can extract, from 2P imaging data, all or most neural information available in the underlying spike trains, we first implemented a realistic forward model for the generation of synthetic fluorescence data from ground truth spike trains. This forward model is available within NIT and can be used by users to perform their own simulated experiments to match their own experimental conditions. We implemented and compared two models for the generation of synthetic two-photon calcium imaging traces.

The first one (**Figure 2A**, left panel) defines the spike to fluorescence transfer function through a linear convolution with a double-exponential kernel [116–118]. This model is a good approximation of the fluorescence evoked by action potentials in a low spike rate regime, but fails to account for non-linear effects present at high firing rates [119].

**Figure 2.**
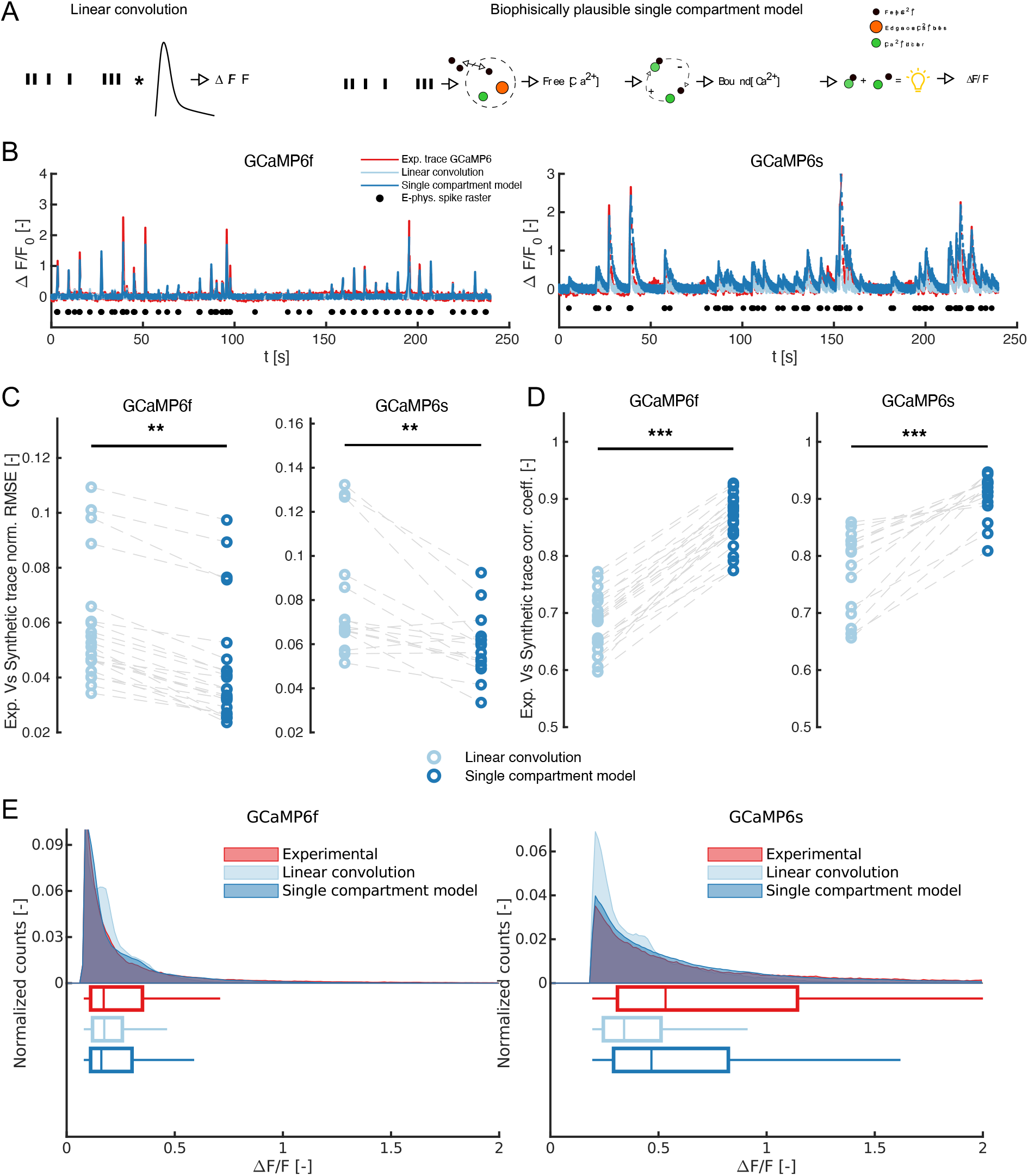
Comparison of methods for the generation of synthetic GCaMP6 traces given a spike train. (**A**) Schematics of the two methods considered: a linear convolution of the spike train with a double exponential kernel (left) and a biophysically plausible Single Compartment Model (SCM) of calcium dynamics (right). The SCM considers the presence of endogenous (orange) and exogenous (green) calcium buffers in the cytoplasm to predict the concentration of free calcium within the cell membrane. Binding/unbinding dynamics of free calcium to the indicator is simulated to generate time traces of bound and un-bound fluorophore concentrations. Synthetic GCaMP6 fluorescence traces are then generated through a linear combination of the concentration of bound and un-bound indicator concentrations. (**B**) Sample two-photon GCaMP6 experimental traces (red) recorded with simultaneous loose-seal cell-attached electrophysiology (black scatter). Experimental data from [122],[121]. The panel also shows synthetic traces generated using both a linear convolution (light blue) and SCM (dark blue) given the experimentally recorded spike train, under the same SNR than the experimental GCaMP6 trace. (**C**) RMSE of synthetic Vs experimental GCaMP6 traces for both models considered (**: p < 0.01, one-tailed Kruskal-Wallis test). (**D**) Correlation coefficient of synthetic Vs experimental GCaMP6 traces for both models considered (***: p < 0.001, one-tailed Kruskal-Wallis test). (**E**) Distribution of the upper 30^th^ percentile of ΔF/F values across all frames in experimental data and both linear convolution and SCM models.

The second model (**Figure 2A**, right panel) is based on a single compartment model (SCM) of calcium dynamics in the cytoplasm [120]. Generation of fluorescence from a given spike train is obtained in three successive steps. The first step models the concentration of unbound calcium within the cell membrane. Every action potential elicits a step influx of calcium ions. The free calcium intake accounts, in a non-linear way, for the effects of both endogenous and exogenous (indicator) calcium buffers in the cytoplasm. The extraction of free calcium from the cell is modelled through a linear leak term combined with a non-linear extrusion term for the membrane calcium pumps. Non-linear effect of the release of free calcium from internal buffers in the cell is also included in the model. A second step in the model allows to calculate the fraction of calcium indicator that is bound to calcium to the one that is not. This is performed through integration of the indicator binding/unbinding kinetics. A linear model converts the fraction of bound and unbound indicator to fluorescence values. This biophysically plausible model for fluorescence generation includes four forms of non-linearity, which cannot be obviously present in the linear convolution model. Those are related to: calcium intake after every action potential, free calcium release from endogenous and exogenous buffers, calcium extraction from membrane pumps and saturation of calcium indicator. A sample train of action potential and the resulting traces for free cytoplasmatic calcium, indicator-bound calcium and fluorescence is shown in **Supplementary figure 1**.

In both models, we added Gaussian white noise to the generated fluorescence to account for experimental noise and manipulate the SNR of simulated recordings (see Methods for details). We assessed the accuracy of the two methods in generating realistic calcium imaging traces by comparing synthetic traces with experimental ones. The experimental dataset we used [121, 122] contains simultaneous calcium imaging t-series and juxtasomal electrophysiological recording in neurons expressing both GCaMP6f and GCaMP6s. We used the experimentally recorded action potentials as inputs for both forward models. The levels of noise in the synthetic traces were tuned so that each synthetic ΔF/F signal had the same signal-to-noise ratio (SNR) than the corresponding experimental trace. The sample experimental and synthetic ΔF/F traces, on both indicators, are reported in (**Figure 2B**).

For each acquisition in the dataset, both Root Mean Square Error (RMSE) (**Figure 2C**) and Pearson’s correlation coefficient (**Figure 2D**) between experimental and synthetic ΔF/F traces were calculated. The single compartment model showed significantly better performance than the linear convolution model, both in terms of RMSE and correlation for both considered calcium indicators. To further compare the performance of the two methods, we assessed their performance in reproducing realistically high levels of fluorescence. To this end, we compared the distribution of synthetic ΔF/F values against real values reported by experimental 2P calcium imaging traces (**Figure 2E**). The SCM shows a longer tail of high ΔF/F values – especially evident for GCaMP6s – which is closer to the distribution of the experimental data. This shows that the SCM model allows to generate synthetic 2P calcium imaging traces covering a broader part of the dynamic range of the indicator with respect to a linear convolution kernel. Overall, these results show that the SCM generates more realistic synthetic calcium imaging traces. Thus, in all subsequent NIT information algorithm testing, we used calcium traces generated with the SCM.

#### Effect of neuronal firing and experimental conditions on information available from calcium imaging traces

Recording somatic calcium concentration in neurons through fluorescent two-photon imaging is widely used to infer the neuronal supra-threshold activity [122–128]. However, we still lack a systematic appreciation of the consequences of the limitations of calcium imaging for information-theoretic measures of neural activity and of how best to deal with them. For this reason, we investigated the effect of a series of variables on calculations of information from 2P calcium imaging traces. These include factors related to the underlying neurobiology, such as the shape of post-stimulus time histogram (PSTH), mean spiking rate (SR) to different stimuli, or technical characteristics of the experimental setup, such as imaging frame rate (FR), signal-to-noise ratio (SNR), calcium indicator. We performed a parametric sweep over those parameters as follows.

We simulated activity in response to two different categorical “stimuli” (the variable *S*, s=1 or s=2, in the MI calculation, Equation (1)). These simulated stimuli elicit a different neuronal response over a 1 second post-stimulus window. Differences in stimuli are modeled as differences in the strength and time pattern of the neural responses they elicit, as explained next. The two stimuli could elicit a time-averaged spike rate (SR) along the trial of either 1 or 2 Hz (we termed those cases as *Low MI, low SR*), 12 Hz and 13 Hz (*Low MI, high SR*) and 2 Hz and 12 Hz (*High MI*). For each mean firing rate response, we considered two different temporal shapes of elicited Post-Stimulus-Time-Histograms (PSTHs): *tonic* (i.e. uniform over time) and *phasic* (i.e. Gaussian-shaped time dependency, peaking at 0.25 s, standard deviation 0.01 s). Given a time-averaged SR, both phasic and tonic responses have the same integral over time, i.e. the same expected number of spikes. The shapes of the PSTH are plotted **Figure 3A**, top panels. Spike trains were generated through an inhomogeneous Poisson process with an instantaneous rate equal to stimulus-evoked PSTH. We simulated situations with three different frame rates for the imaging set-up: 5 Hz (representative of galvanometric imaging with raster scanning), 30 Hz (representative of imaging with resonant scanners) and 100 Hz (representative of alternative high acquisition frequency methods, e.g., smart line scanning imaging [126]). Spike trains and ΔF/F traces were always generated at a sampling rate of 1 kHz, and the latter were then subsampled to the desired sampling rate. SNR was varied systematically across simulations by varying the amplitude of the noise added to the calcium imaging traces.

**Figure 3.**
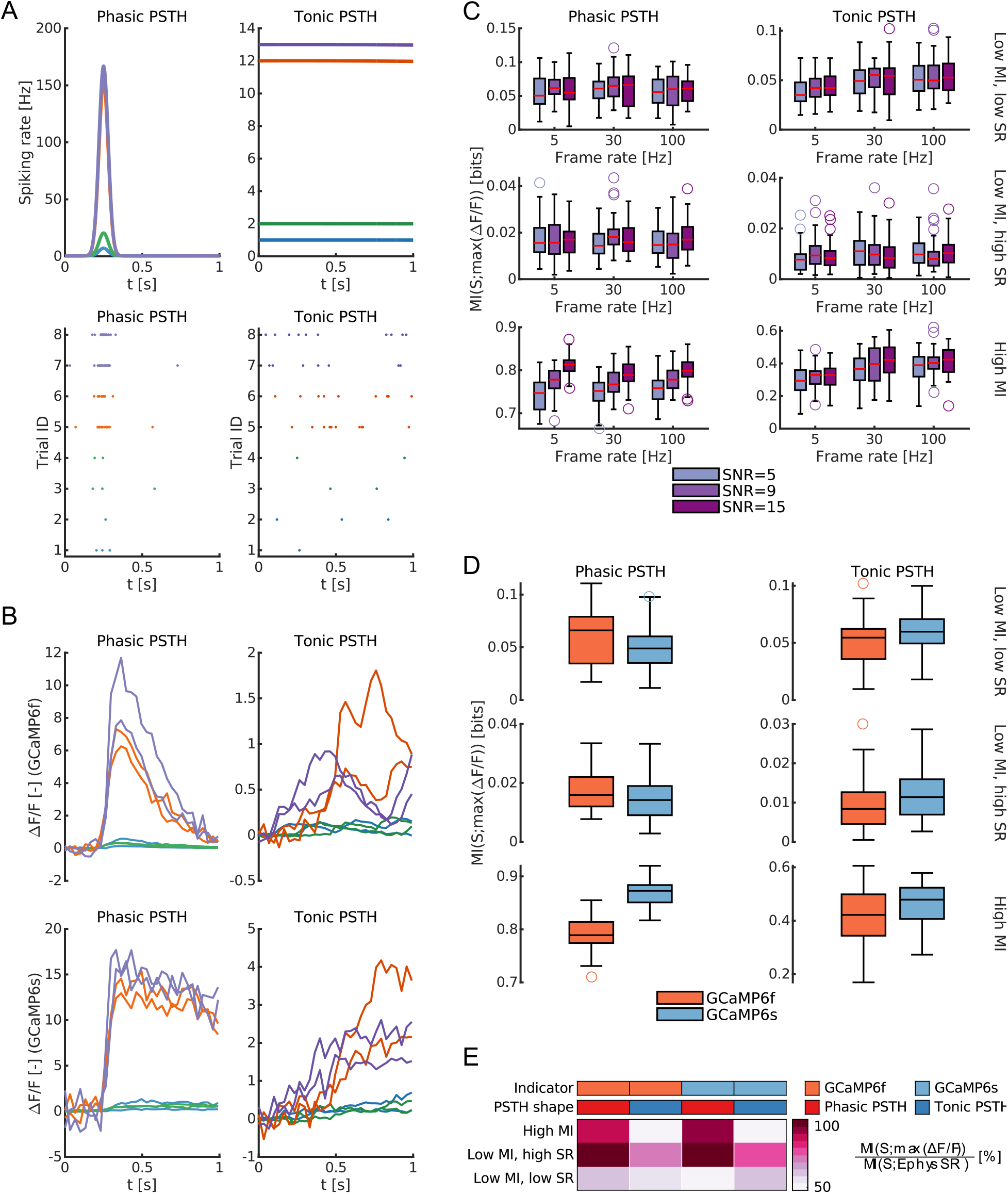
Effect of neuronal firing regime and experimental conditions on stimulus information retrieved from calcium imaging signals. (**A**) Instantaneous neuron spiking rate (SR) for phasic and tonic post-stimulus time histogram (PSTH) responses (top row), average firing rates over the trial duration are identical between the two conditions at fixed stimulus. Corresponding Poisson spike rasters for two sample trials per each stimulus (bottom row). (**B**) Synthetic GCaMP6f (top row) and GCaMP6s (bottom row) traces (SNR 9, frame rate 30 Hz) relative to spike rasters in panel (**A**). (**C**) Distributions of stimulus information in GCaMP6f ΔF/F traces at various information levels and for both tonic and phasic PSTH. Effect of SNR and imaging frame rate on stimulus information. All calculations of MI consider two stimuli. In *Low MI, low SR* the neuron responds to the two stimuli with 1 Hz and 2 Hz average spiking rate (blue and green curves panel A). In *Low MI, high SR* the neuron responds to the two stimuli with 12 Hz and 13 Hz average spiking rate (orange and violet curves panel A). In *High MI* the neuron responds to the two stimuli with 2 Hz and 12 Hz average spiking rate (green and orange curves panel A). Each box plot reports data from 50 simulations. Results of the statistical analysis for the data in this panel are reported in **Supplementary table 5**. (**D**) Effect of calcium indicator on stimulus information at different PSTH shapes and information levels. Each box plot reports data from 50 simulations. Results of the statistical analysis for the data in this panel are reported in **Supplementary table 6**. (**E**) Percent of stimulus information in max ΔF/F with respect to MI encoded in spike rate at the same conditions. Values are average values over 50 simulations. All data in the figure refer to simulated traces. Mutual information is evaluated using plug-in method. All MI calculations consider max ΔF/F across the trial as a metric of neuronal response.

Sample spike trains and ΔF/F traces (30 Hz frame rate, SNR = 15, two sample trials per each mean firing rate) for both GCaMP6f and GCaMP6s are shown in **Figure 3B**. In this part of the analysis, informaiton calculation parameters were as follows. We used the plug-in direct method, discretizing these neural responses in 4 equi-spaced bins. We used peak ΔF/F over the trial as response *R*, as it is a widely used approach for the analysis of two-photon imaging data [64, 129]. For each combination of parameters (SNR, FR, calcium indicator, PSTH and levels of stimulus-modulated firing rate), 50 independent MI calculations (each with 400 trials per stimulus) were performed. No limited sampling bias correction was used, because the number of trials was large enough for the MI to be bias-free [14].

We first investigated the effect of varying the imaging FR and SNR on the mutual information computed from the somatic calcium imaging signal for phasic and tonic PSTH shapes (**Figure 3C**, results of the statistical tests are summarized in **Supplementary table 5**). In **Figure 3C** we used peak ΔF/F of GCaMP6f to compute information from the calcium traces, but we obtained similar results (not shown) using other calcium imaging metrics (e.g. mean ΔF/F). Both FR and SNR have a limited effect size on the information contained in the peak ΔF/F. The notable exception was the case of phasic PSTH shapes and high neural information, in which case increasing SNR led to a notable increase of stimulus information with SNR (**Figure 3C** and **Supplementary table 5**). The effect of using either a slower (GCaMP6s) or faster (GCaMP6f) calcium indicator is explored in **Figure 3D** and **Supplementary table 6** (with SNR = 15 frame rate = 30 Hz). In most cases the information obtained from the calcium traces with peak ΔF/F was approximately the same with either indicator, with the exception of the high information, phasic PSTH case. In this case using the GCaMP6s led to higher information extracted from the calcium traces, due to its slower dynamics and higher dynamic range compared to GCaMP6f.

Because calcium imaging measures indirectly the neural activity, with a lower SNR and lower temporal resolution that direct electrophysiological recording of spikes, it is commonly assumed that the information reported by a calcium indicator will be smaller than that encoded in neural activity. To evaluate this information loss we computed, the average fraction of information present in peak ΔF/F, relative to the one present in a spike rate code. We found that the percentage of spike rate information extracted on average from the calcium traces varied widely, from 50% to 100% (**Figure 3E**), depending in particular on the features of neuronal firing. More stimulus information is lost when computing it from the calcium traces rather than from the spike rate when the simulated neuron fires tonically than when it fires in a phasic way. This is because, as apparent from the individual traces in **Figure 3B**, the phasic PSTHs with a stronger and more concentrated spike rate elicit more repeatable and less noisy calcium traces than those obtained with the tonic PSTHs having a similar number of spikes randomly distributed over time.

In sum, our simulations suggest that the NIT information theoretic analysis of calcium traces recovers a good fraction (between 50% and 100%) of the information encoded in electrophysiological spike rates, with the extraction being particularly efficient for high-rate phasic responses and high dynamic range indicators.

#### Spike rate information is not an upper bound for stimulus information contained in ΔF/F traces

Since as discussed above calcium imaging reports an indirect measure of neural spiking activity, the information about stimuli computed from ΔF/F traces will miss out on some of the information carried by the temporal spike pattern as measured from electrophysiology recordings. However, this does not necessarily imply that in all cases the information computed from the calcium traces will be lower than the information carried by the underlying spike rate code.

From the mathematical point of view, the data processing inequality [98] ensures that stimulus information cannot be increased, but can only be lost or remain equal, every time a transformation of *R* not dependent on *S* is applied to the data. This implies that information in the spike rate is always lower than or equal to the information contained in the full spike train. However, because the transformation that maps the spike train into a calcium trace is not a direct consequence, in Markovian terms, of the transformation that links a spike train to spike rate, the stimulus information in the calcium trace may either be higher, equal or lower than the stimulus information in a rate code.

From the intuitive neurobiological point of view, the fluorescence traces can have more information that the spike rate in cases in which the latter loses some of the information encoded in the spike timing that the former captures. Indeed, owing to the slow dynamics of the indicator, ΔF/F traces contain not only information about how many spikes are emitted by a neuron, but also how close they are in time. The contribution of this effect to the information content of calcium traces is amplified as the ratio between the decay constant of the indicator and the stimulus-modulated inter-spike interval increases, and as the informative content of a spike-rate code alone decreases. As such, it becomes particularly evident for phasic PSTH when stimulus information is encoded at high mean firing rates and rate information is low (**Figure 4A**). Data in **Figure 4** are from a limited portion of the full parametric sweep (FR = 5 Hz, SNR = 15, GCaMP6f), but similar conclusions can be drawn when considering the full range of parameters investigated (**Supplementary figure 4**). As an example, we have considered one of the points (green scatter in **Figure 4A**, central panel) showing more information in peak ΔF/F than in SR.

**Figure 4.**
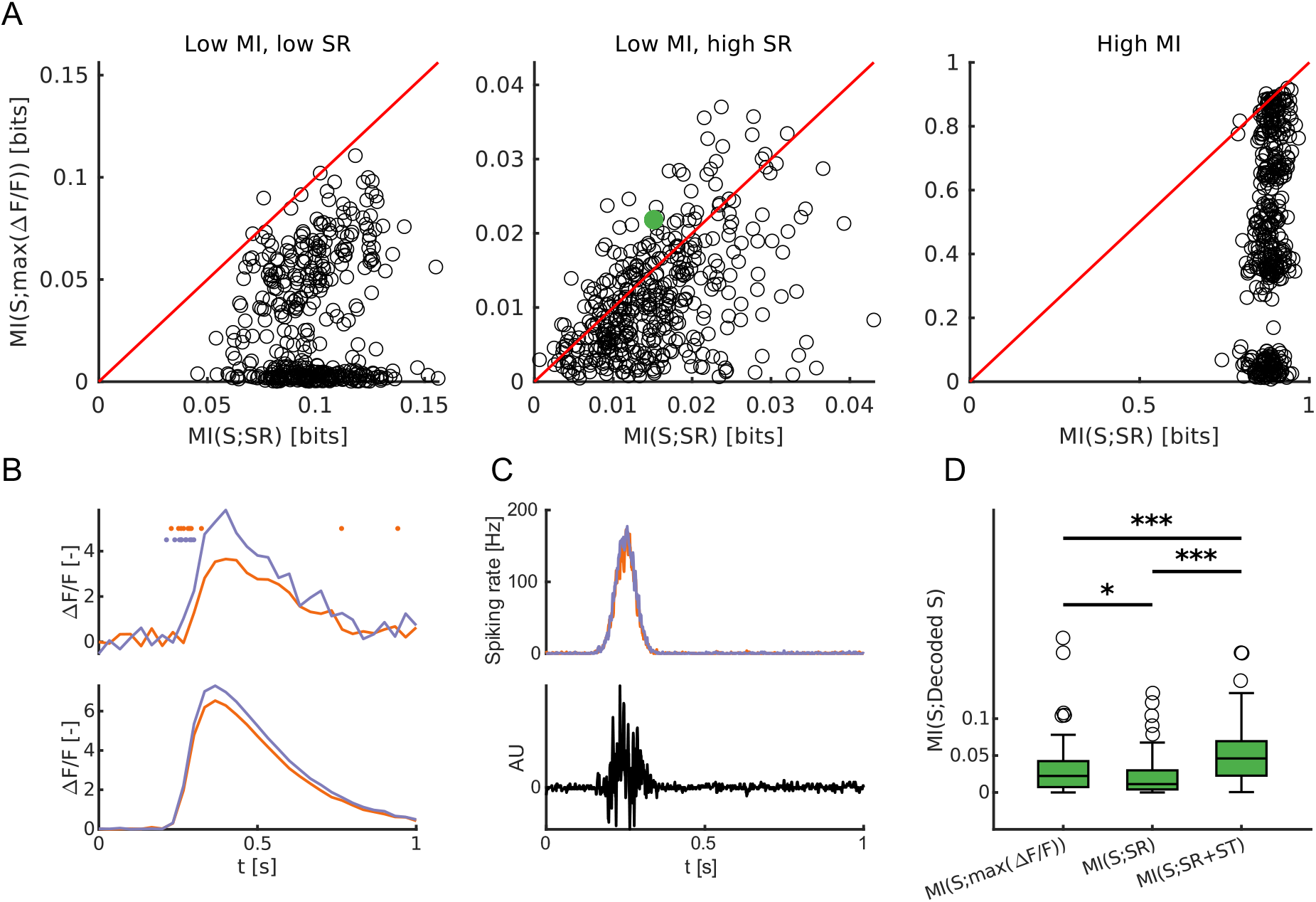
Information in ΔF/F can be higher than spike rate information. (**A**) Scatter plots of stimulus information in SR vs stimulus information in peak ΔF/F. Each scatter results from one over 50 MI calculations across the following parametric sweep: SNR = 15, FR = 30 Hz, GCaMP6f, phasic PSTH. Red lines are the quadrant bisectors. The green scatter point refers to the point analyzed in panels (**B**-**D**). (**B**) Top: stimulus-evoked spike rasters and corresponding ΔF/F traces for two specific trials with an identical trial-averaged spike rate (10 Hz) but responding to two different stimuli (color-coded). Bottom: trial-averaged stimulus-evoked ΔF/F traces. (**C**) Top: trial-averaged PSTH for the response to the two stimuli. Bottom: spike-timing template used in the decoding analysis in panel (**D**). (**D**) Values of MI between true and decoded stimulus calculated when considering: max ΔF/F, SR and simultaneous contribution of SR and spike timing (ST). The analysis is performed on the data corresponding to the green point in panel (**A**). Box plots report 100 cross-validated runs of GLM decoder (*: p < 0.05, ***: p < 0.001, Bonferroni corrected Kruskal-Wallis multiple comparison test). All data in the figure refer to simulated traces. Mutual information is evaluated using plug-in method.

In this case, because of the different stimulus-modulated inter-spike interval, even when the two stimuli elicit an identical spike rate in two different trials, the ΔF/F traces will still show stimulus-related differences (e.g. different peak activity as show in **Figure 4B**, top row) similar to their trial-averages (**Figure 4B**, bottom row). Additionally, for the case of a phasic PSTH, the only stimulus informative spikes are time located in the narrow window around the peak of phasic activity (**Figure 4C**). All other spikes emitted in the baseline activity period (baseline firing rate set at 0.5 Hz in all simulations reported) are non-informative and thus degrade the SR information. On the contrary, given the high stimulus-modulated firing rate of the neurons, and the slow dynamics of calcium indicators, spikes outside of the stimulus-modulated window have little effect on ΔF/F traces, contributing to increase its stimulus information compared to a spike rate code.

Thus, an ideal decoder of neural activity would use the spike times to consider only those spikes in the informative window and discard the others, together with weighting spikes in the informative window proportionally to the instantaneous inter-spike interval. We implemented such decoder by projecting the neural activity in each trial on a template based on the difference between the trial averaged PSTH when responding to the two stimuli (**Figure 4C**, bottom). We have then used the GLM decoder implemented in NIT to calculate the MI between the real and decoded stimulus when using peak ΔF/F, spike rate (SR) or the template projected activity and spike rate (SR+ST), to compare their information content. Results are summarized in **Figure 4D**. Each box plot in the figure shows the distributions of MI between the real stimulus and the decoded one across 100 cross-validated runs of the GLM classifier. While the stimulus information in the calcium trace is lower than the one present when considering both spike rate and spike timing, it is significantly higher than the mere SR information. This shows how the calcium dynamics captures some properties of the optimal spike timing decoder and that spike timing contributes to the informative content represented in ΔF/F.

While cases like the above example – in which more information is available in the calcium traces than in the time-averaged spike rates – may not happen frequently with real data, it should be noted that calcium traces will always contain a mix of spike rate and spike timing information, which is important to keep in mind when interpreting empirical results.

#### Dependence of stimulus information on the metric used to quantify single-trial calcium fluorescence responses

In the previous sections, we quantified information from calcium traces using the peak ΔF/F as a metric of single-trial responses based on two-photon fluorescence. This measure is widely used in the analysis of calcium imaging data [64, 129–131], but is not the only possible choice. Several other metrics are commonly used to quantify single-trial activity in a post-stimulus window from calcium imaging signals. These metrics include: mean ΔF/F [64], integral ΔF/F [132–135], linear deconvolution using an exponential kernel [136], spike inference algorithms [127, 137–139]. Among spike inference methods, we focused on OASIS [137] due its competitive performance [117].

To inform future information-theoretic analyses of calcium imaging traces, we investigated on simulated data how well the different metrics listed above performed in extracting stimulus information. All listed metrics have advantages and disadvantages. The peak ΔF/F captures the strength of the calcium transient responses but can be heavily influenced by noise and does not capture the temporal structure of the fluorescence. Mean and integral ΔF/F are less influenced by noise, but they are less effective in capturing the strength of transient activations. Both linear deconvolution and OASIS quantify aspects of calcium signal potentially closer to spiking activity but assume a linear relation between spikes and measured fluorescence. In addition to the methods listed above, we propose a novel non-linear metric, that we termed *estimated calcium*, that inverts the biophysically plausible non-linear forward model to estimate the concentration of intracellular free calcium from ΔF/F traces (see Materials and methods).

We have thus used the same five-dimensional sweep of simulation parameters (FR, SNR, indicator, PSTH shape and stimulus modulation of SR) used in **Figure 3** to calculate the levels of stimulus information contained in each of the above-mentioned measures of neural activity in the 1-second-long post-stimulus window. We computed stimulus MI in both SR and ΔF/F metrics using the direct method with equally-spaced binning in 4 bins. Fifty independent runs are performed in each of the coordinate points of the parametric sweep. The distribution of ΔF/F metrics showing the highest mean amount of stimulus information across the parametric sweep is shown in **Figure 5A** (actual levels of MI across all conditions in the parametric sweep are reported in **Figure 5B**, together with the value of stimulus information in the spike rate code). Overall, peak ΔF/F extracts most stimulus information when the stimulus is encoded at high rates, mostly when the neuronal response has a tonic PSTH. In these conditions the stimulus will, in fact, modulate mostly the amplitude of the calcium imaging response. In other conditions, most of the stimulus information contained in the calcium imaging response was retrieved by estimated calcium. OASIS shows good performance at high imaging frame rates, though it suffers particularly low rates (**Supplementary figure 3**). When looking at the absolute levels of information retrieved across all conditions **Figure 5B**, estimated calcium performs on average better than the other metrics considered by recovering about 65% of the underlying SR code.

**Figure 5.**
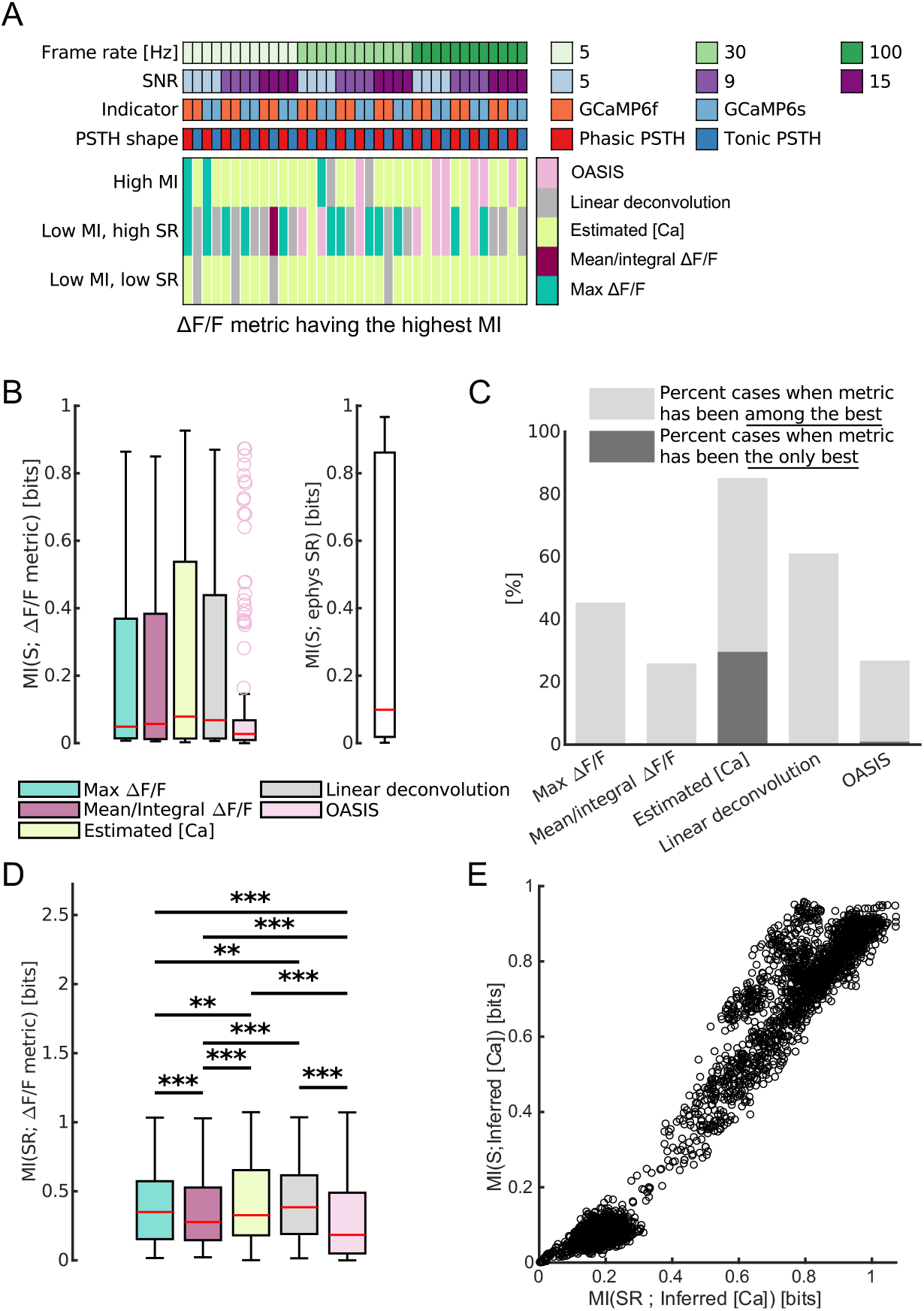
Appropriate processing of ΔF/F signal increases the retrieved stimulus information from calcium imaging traces. (**A**) Best performing metric based on ΔF/F signal for each of the conditions explored in the parametric sweep. Best performing metric at each condition is defined as the one retrieving the highest value of stimulus information. See Materials and methods for detailed definitions of each of the metrics. (**B**) Distributions of values of stimulus information reported by each metric (left) and by a spike rate code (right) across all the calculations performed in the parametric sweep shown in panel (A). (**C**) Percent of cases, across the whole parametric sweep shown in panel (A), where each ΔF/F metric has been among the best performing metrics (light grey bars) or the single best performing one (dark grey bars). Best metrics are defined as the ones recovering the highest amount of stimulus information (p < 0.05 Bonferroni corrected Kruskal-Wallis multiple comparison test). Full data for this figure are reported in Supplementary figure 3. (**D**) Distributions of values of spike rate information reported by each metric across all the calculations performed in the parametric sweep shown in panel (A). (**E**) Scatter plot of stimulus information in Inferred [Ca] against spike rate information carried by the same metric. Data in this panel include All data in the figure refer to simulated traces. Mutual information is evaluated using plug-in method using 4 equally-spaced bins do discretize spike rate and the calcium metrics.

Statistical significance of the results was assessed through Kruskal-Wallis test with Bonferroni correction for post-hoc comparisons. In all conditions of the parametric sweep, the best performing calcium imaging metric, together with the others being non statistically different from it (p > 0.05), were marked as best for that condition (stars in **Supplementary figure 3**). **Figure 5C** reports the percentage of cases, across all the conditions examined, where each metric was part (light grey) – or was the only component (dark grey) - of the best performing group. Both the linear deconvolution and the newly proposed estimated [Ca] showed to be the most versatile spiking activity metrics based on ΔF/F. Estimated [Ca] is among the best performing metrics in more than 80% of the conditions examined in our parametric sweep and is the only best performing one in around 25% of cases considered. Linear deconvolution works well in retrieving stimulus information in around 60% of conditions. Mean/integral ΔF/F and OASIS, on the other side, are only among the best performing groups in around 25% of the cases, showing poorer performance in reconstructing spiking information. (Note that the poorer performance of OASIS was not due to incorrect set-up of the algorithm as we have verified (**Supplementary figure 2**) that the deconvolved activity we obtained through OASIS had similar correlation with ground-truth spike recordings as reported previously using this algorithm on the same dataset we use here for validation, see [140]). Linear regression of the average z-scored deconvolved activity using OASIS and the underlying ground truth SR shows, however, how the levels of z-scored deconvolved activity predicted by OASIS have a relatively high variability that cannot be explained by a linear fit (R2 = 0.52 GCaMP6f, R2 = 0.34 GCaMP6s). This suggests that, while OASIS matches the timing of neuronal activity with reasonable accuracy, the magnitude of the deconvolved calcium trace reflects less well the underlying firing rate, limiting the applicability of the method for information-theoretic measures of neuronal activity. The poorer performance of OASIS becomes especially noteworthy given that spike inference algorithms are typically performing better on synthetic data than in real experimental conditions, and the assumption of Poisson spiking used in our synthetic data should favor the method’s accuracy [137].

In addition to considering which calcium metrics are better for computing single-trial information about external stimuli, we consider another, and perhaps equally important question, of which metric of calcium activity best reconstructs the underlying spike rate of the same cell. We computed the mutual information between the spike rate during the 1s post-stimulus window in our simulated trials and the calcium metric. This result (**Figure 5D**) confirms that estimated calcium and a linear deconvolution are, on average, carrying more information about the spike rate code than the other calcium metrics analyzed.

An explanation of why calcium metrics that carry higher stimulus information also carry higher information about the spike rate is that stimulus information is carried by the spiking activity of neurons and these calcium metrics reconstruct its value well. In support of this explanation, we found that the levels of stimulus information extracted from the ΔF/F activity with a given set of simulation parameters correlated with the levels of information present between the calcium imaging signal and the electrophysiology in the same simulation, as shown by the scatterplot of the two information values across different simulations for the case of the estimated calcium (**Figure 5E**).

#### A comparison of non-parametric copula and binned plug-in methods for computing information from calcium imaging traces

All above examples computed information using the plug-in binned methods, a choice that has been widely used due to its ease of implementation, robustness and fast computational time [25, 27, 38, 54]. However, other more computational demanding but potentially more accurate methods are also available to compute information from limited experimental samples. NIT implements the recently developed Non-Parametric-Copula information estimation [71]. Here we test the advantages for computation of information from calcium imaging of this more computationally expensive method.

We first investigated whether the NPC offers advantages in terms of reduction of limited sampling bias in case limited datasets are available. To this end, we introduced, in the multidimensional sweep over the simulation parameter space outlined in the previous sections, a further parameter: the available number of trials per stimulus (here varied in the range 5 to 400). We found (**Figure 6A**) that, for both copula and direct plugin method, and consistent with previous studies [14], the information had a big upward bias for low numbers of trial per stimulus (5 to 20), and then converged to the asymptotic value for larger number of trials (several tens). To quantify how quickly the information estimate in individual simulations reached the asymptotic values across methods, we repeated the above analysis over a large number of simulations with different parameters according to our 5-dimensional parameter sweep. For each individual set of simulation parameters, we compared the distribution of calcium information values for different numbers of trials against the asymptotic (400 trials per stimulus) distribution. The lowest number of trials giving a distribution not significantly different from asymptotic (t-test, p-val < 0.05) was considered the minimum required by the method to provide a bias-free estimate of MI. We repeated the process for the whole parametric sweep, computing the ratio between the trials needed by the copula and by the binned methods. The distribution of the ratio is shown in **Figure 6B**. In this figure, values lower than 1 imply that the copula method is performing better than binned methods for bias free information estimations, while values higher than 1 imply than the binning method works better. For the vast majority of simulations, the non-parametric copula needed less trials to reach asymptotic values of information. Thus, the non-parametric copula should be favored when analyzing smaller datasets.

**Figure 6.**
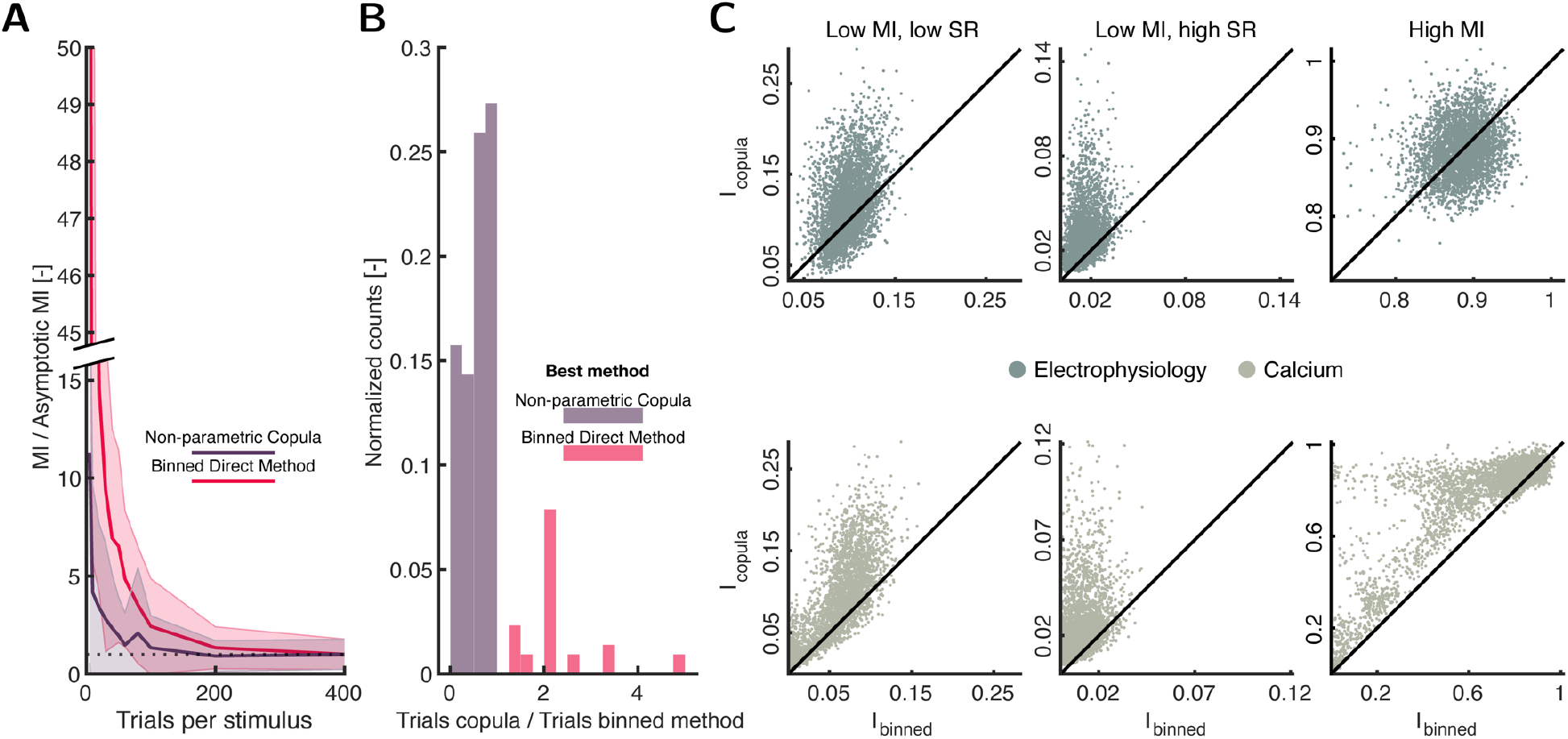
Comparison between binned methods and non-parametric copula. (**A**) MI values (mean±SD) for a single coordinate point in the considered parametric sweep (GCaMP6f, Frame Rate = 5 Hz, SNR = 5, *Low MI, High SR*, tonic PSTH) using NPC and binned direct method with an increasing number of trials per stimulus. The dotted horizonal line represents the y-axis value of one (when the information estimation reaches its asymptotic value) (**B**) Distribution of ratio between number of trials needed by the copula and binned method to reach asymptotic information values. Note that a ratio lower than 1 implies that the copula retrieves asymptotic values with less trials than binned method. (**C**) Information values provided by the copula against values given by the binned method for three different information levels (*Low MI, low spike rate; Mid MI, high spike rate;* and *High MI*) and both electrophysiology and calcium data. All data in the figure refer to simulated traces. Note that the samples included in this figure correspond only to the deconvolved calcium.

Non-parametric copula is particularly suited to be applied on continuous variables. This suggests that larger amounts of information can be extracted from calcium signals, which are continuous, with non-parametric copulas than with binned estimators for MI. We thus examined the asymptotic information values provided by the copula against binned methods for both simulated spike trains and calcium imaging traces (**Figure 6C**). For the electrophysiology, the non-parametric copula performed better than binned methods only in the cases which information content is not high. However, for calcium imaging, the advantage of the copula was accentuated and was also present in high information cases. This underlies the specify usefulness of the non-parametric copula for calcium imaging. Note that we did not find comparably high performance when using parametric Gaussian copulas (also implemented in NIT), rather than non-parametric copulas. This is because responses of individual neurons carry non-gaussian dependencies with the stimulus (statistics is of neurons approximately Poisson, which differs from Gaussians for low spike numbers typically observed in a trial) and this translates in non-gaussian dependence between stimuli and calcium traces, which make the use of Gaussian copulas not generally applicable.

Despite the advantages that the copula has compared to the binned methods, there also exist drawbacks main limitation is the computational time required to fit the copula based on the data. As an example, the computations reported in **Figure 6A** required approximatively 200 time more CPU time with the non-parametric copula than the direct plug-in method.

### Analysis of experimental data validates findings on synthetic traces

Our information theoretic analysis of realistic simulations of calcium imaging traces generated by neural spiking activity indicates that the calcium imaging traces are able to extract sizeable amounts information about both external stimuli and about the levels of the underlying spike rates. It also suggests that certain metrics of single-trial activity for calcium traces are better than others for extracting such information. Here, we tested some of the above predictions from simulated activity on real empirical data. We used NIT to analyze four independent datasets with simultaneous cell-attached electrophysiological and two-photon imaging recordings from both GCaMP6f and GCaMP6s-labelled neurons during spontaneous activity [119, 122, 141, 142]. We focused on using NIT to compute how much information about the spike rate each calcium metric provides. We divided the experimental time traces in padded windows of 0.5s, and then computed the mutual information between the spike rate in a considered window and the calcium metric in the given window. We used the NPC information calculation method as it performs more reliably as shown in the previous section. Similar conclusions, however, would have been reached using the direct binned method (not shown).

Results of this information calculation on all neurons with calcium traces with SNR higher than 9 are reported in **Figure 7.** These results confirm that, as with the simulated data, sizeable amount of information about the underlying spike rate can be obtained from the underlying traces. These information values are of the order of 0.2 bits, which corresponds with significant but far from perfect spike train reconstruction from the calcium metrics. Comparison of how the amount of information varies between ΔF/F (**Figure 7**) confirms the results emerging from the parametric sweep on simulated traces. Estimated calcium and linear deconvolution were, on average, better at reconstructing spike train information that other calcium imaging metrics.

**Figure 7.**
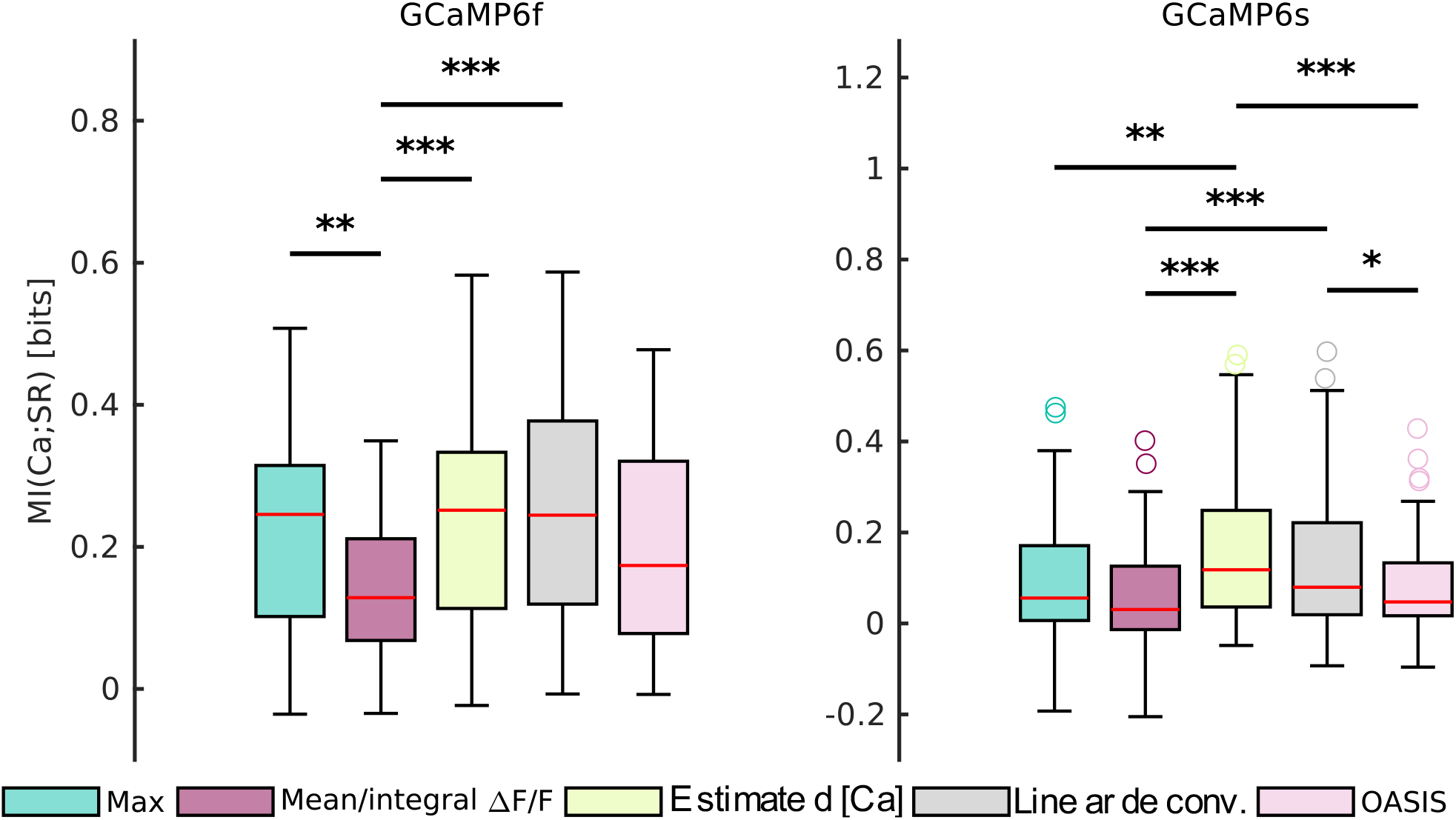
Validation of performance of spiking activity metrics based on ΔF/F in recovering stimulus information on experimental data. Box plots of mutual information between different spiking activity metrics based on ΔF/F and spike rate. Data in this panel refer to simultaneous cell-attached electrophysiology and two-photon imaging recordings from previous publications [119, 122, 141, 143]. Traces in the original datasets have been filtered for SNR > 9.

The publicly available datasets were designed to test the correspondence between spike rates and calcium traces and not to study sensory coding, thus had no or insufficient data with responses to sensory stimuli to study sensory information. However, as shown by our simulations (**Figure 5E**), metrics that are appropriate for inferring spike rate values are also expected to be appropriate to extract stimulus information.

### Examples of use of intersection information to find pure, stimulus unrelated choice signals as markers of preparatory activity

We finally exemplify, on real data, possible uses of the PID tools within NIT. In particularly, we exemplify possible and novel uses of Intersection Information (II) [82, 83], a formalism developed specifically for the analysis of neural recordings in perceptual decision tasks. As reviewed in the Intersection Information section above, II measures that amount of information carried by neural activity that is shared by both stimulus and choice. Thus, II can be interpreted as the part of stimulus information carried by neural activity that is also choice information. In this interpretation, II has been applied to sensory neuron to investigate the extent to which the information encoded in sensory areas is relevant to form behavioral choices [39, 54, 83]. For example, it has been used to investigate whether in primary and secondary somatosensory cortices the behavioral discrimination of texture of surfaces is supported by the texture information encoded in millisecond-precise spike times or in spike rates [83, 144]. The authors found that on average similar amounts of texture information were encoded by the millisecond precise spike times and by the spike rate of neurons. However, the behavioral discrimination performance of the rat was higher when spike times provided correct texture information than when spike times provided incorrect information, whereas behavioral performance did not depend much on the correctness of the information provided in spike rates [144]. As a consequence, the amount II carried by spike times was 3 times larger than that carried by spike rates [83], demonstrating that the texture information carried by spike timing has a much larger impact on forming correct behavioral choices than the information carried by spike rates. This type of reasoning is helpful informing hypotheses about the neural code used for sensory perception [82], as it takes into account not only the amount of information encoded in neural activity but also its impact on trial-to-trial behavioral discriminations.

Here, to demonstrate the usefulness of this approach also in contexts different from sensory perception, we use II implemented in NIT to uncover the presence of preparatory motor activity in motor cortices. We applied NIT to a publicly available dataset [145] of 2P calcium-imaging recordings in anterolateral (ALM) and medial motor (MM) cortex of Thy1-GCaMP6s transgenic mice collected during a tactile delayed two alternative forced choice (2AFC) discrimination task (see **Figure 8A** and **B**). Mice were trained to discriminate a pole in an anterior or posterior location using their whiskers. The stimulus was presented for 1.2 seconds during the Sample epoch, followed by a Delay epoch of 3s for the mice to plan the action. A Go Cue indicated the Response epoch for mice to report their guess. In the original publication [145], the authors analysis these recordings with a 3-way ANOVA, including as factors selectivity to the sensory stimulus, the choice reported by the animal, and the trial outcome (correct vs incorrect discrimination). The authors found earlier choice signal in ALM than in MM, suggesting therefore that preparatory motor activity arises first in ALM than in MM. The ANOVA analysis does not include non-linear tuning effects, and does not per se provide a quantification of the values available for single trial discrimination. These issues can be better addressed with information theory. We first computed, using Shannon Information (Equation (1)), the amount of stimulus and choice information carried by the activity of each neuron in short time windows (1 imaging frame, 70 ms) as function of time during the task. Such information values, averaged over all neurons imaged in each area, are reported in **Figure 8C**. We were particularly interested in signals at the beginning of the trial, because they inform more about preparatory activity. In the initial part of the trial (the end of the sample period and the early delay phase), neurons in both areas carried information about both stimulus and choice, with comparable values of stimulus and choice information in ALM and much higher values of stimulus information in MM. Neural activity related to movement preparation can be identified as an early genuinely choice-selective neural signal. However, given that choice and stimulus in each trial are correlated (because the animal performed the task 74% correct, it is possible to predict choice from stimulus), the presence of choice information in neural activity may reflect in full or in part the fact that neurons are actually selective to the stimulus and this in turns make neurons choice selective. To establish the presence of preparatory activity it is thus important to compute presence of pure choice information that cannot be explained by the tuning of stimuli. The formalism of II allows a principled and powerful definition of such pure choice information. II, as explained above, quantifies the amount of information carried by neural activity that is shared by both stimulus and choice. Thus, it quantifies the part of choice information carried by neural activity that is also stimulus information. As a consequence, the difference between II and choice information can be taken as a pure choice information measure, that is a measure of the amount of choice information in neural activity that cannot be explained by the tuning of neurons to stimulus. **Figure 8D** plots the time course of the average amount of instantaneous pure choice information carried on average by the activity of a neuron in a short time window. These results show that, compatible with the results of [145], the pure choice information is present (that is, larger than zero) at approximately 2 s after the pole removal in ALM, but it is not present until 2 seconds later (end of delay epoch) in MM. These results confirm those reported by [145] in a new way that also incorporates the effect of possible non-linearities of tuning of individual neurons.

**Figure 8.**
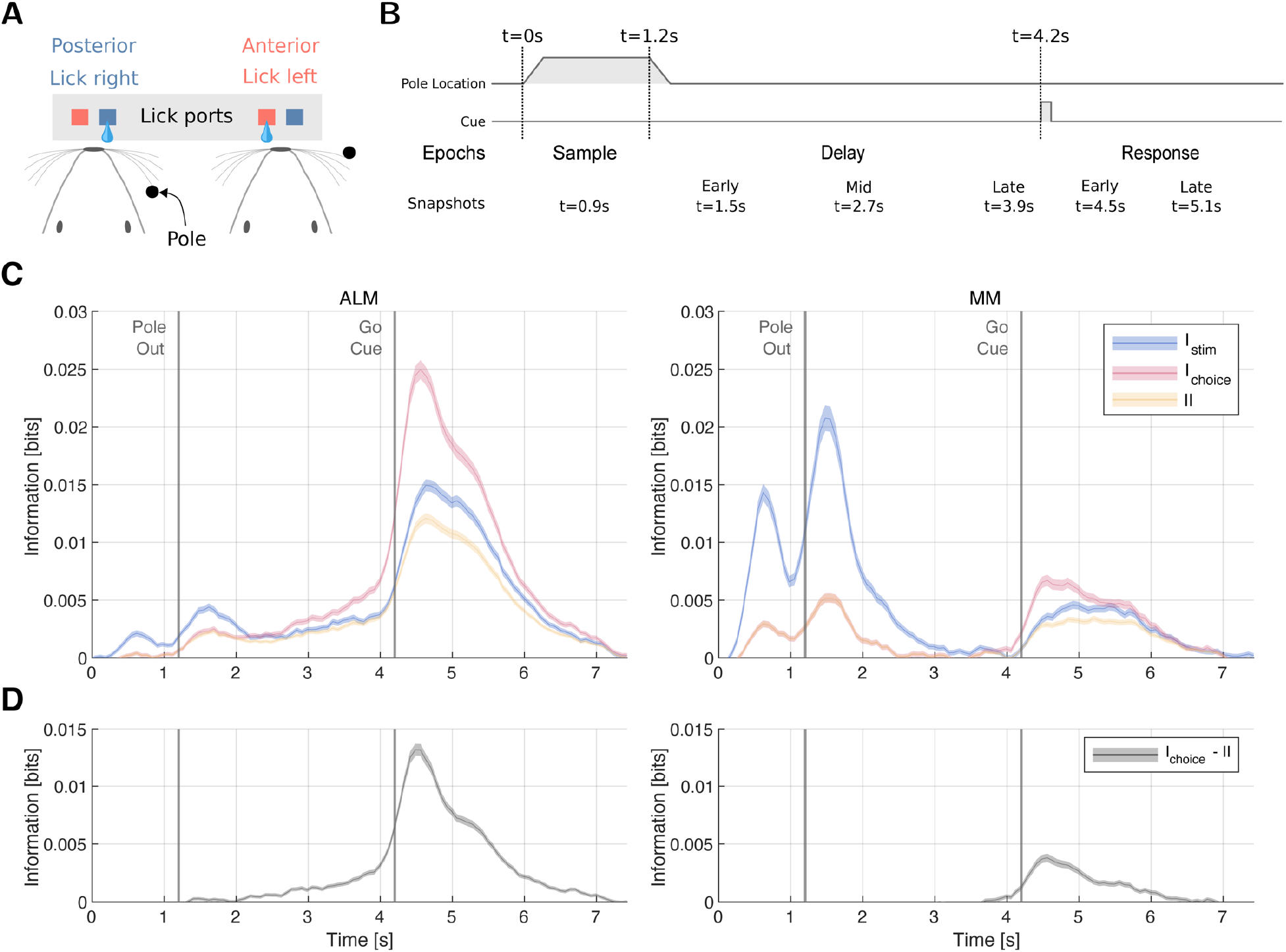
(**A**) Sketch of task. Mice had to lick the right port when the pole was in a posterior location while when in an anterior location, they had to lick the left pole. (**B**) Trial was structured into three different epochs. During the sample epoch (1.2 seconds), the stimulus was provided to the mice. A subsequent delay epoch (3 seconds) without stimulus preceded the Go Cue auditory signal, that initiates the response epoch, in which the mice must report by licking. (**C**) Stimulus, choice, and intersection information over time averaged across neurons computed using 2 bins. Values are bias corrected. Note that we estimated the bias using the average information values found in the pre-stimulus window. (**D**) The difference between choice and intersection information is reported as a proxy of pure choice information measure. Panels A-B redrawn from Ref [145].

## Discussion

The high relevance of information theory for the analysis of neural data calls for open-source, comprehensive, and well documented software packages tailored for neuroscience applications. Here we provide a new such toolbox, NIT, constructed to meet the requirements of the contemporary systems-level neuroscience community. In what follows, we discuss the specific advances of NIT with respect to existing toolboxes and the implications and relevance of our work for neuroscience.

### The breadth of algorithms implemented in NIT can address timely questions in systems neuroscience

Analysis of activity of populations of neurons recorded during the presentation of sensory stimuli and/or performance of cognitive tasks is central to the study of neural coding. Over the last decade, the emphasis of neural coding has shifted from considering purely encoding of sensory information to studying how the encoded information informs choices and behavior [82]. Other prominent current area of investigation include the study of the transmission of information between different brain areas, and the investigation how functions of the brain emerge from interactions among neurons in larger and larger populations [35]. Compared to current information toolboxes, our toolbox adds several important elements to tackle these problems.

NIT supports research on the relevance of neural activity to inform behavioral choices by implementing measures of Intersection Information (II) [82, 83]. II has been proposed and used principally as a measure of how much of the sensory information encoded in neural activity is used to inform choices [39, 54, 82, 83]. This has led to redefine the concept of neural code as the set of features not only carrying sensory information, but also used to drive appropriate behavior [82]. Here, in our application to calcium imaging data (**Figure 8**), we showed how II can be used to address more questions about neural coding than originally proposed. We showed how II can be used to individuate pure choice signals which are not related to stimulus coding. This is of importance in tasks in which sensory signals are associated with the request to executed specific motor programs, such as turning or licking in a certain direction upon the presentation of a certain sensory stimulus.

NIT supports research on transmission of information across areas by implementing directed measures of information transfer, including both Transfer Entropy and Directed Information [84, 85] and it allows the computation of more refined recent measures based on PID [90, 91].

NIT supports research on the emergent properties of population codes by implementing tools that quantify the role of correlations in population codes for creating redundancies and synergies, such as those based on interaction information and the information breakdown [19, 74, 75] and those based on PID [49, 146]. Moreover, NIT implements tools that make analyses scalable to large populations, including unsupervised and supervised advanced dimensionality reduction tools, such as regularized GLM classifiers [39, 99, 100], regularized SVM classifiers [102, 105], and space-by-time Non-Negative Matrix Factorization [107, 147]).

Our public, open source, release of the full NIT code will also contribute to the broad effort towards more effective and reproducible neuroscience, through standardization of tools and methods [148] of which open source analysis software is a core component [149, 150]. In this respect, the integration of NIT with other well established analysis pipelines is facilitated by the MATLAB front-end, which can be directly interfaced with Python through the MATLAB Engine API for Python.

### Comparisons with existing information theoretic toolboxes for neuroscience

The breadth of use of information theory in neuroscience have been supported by several excellent and impactful toolboxes. It is thus of interest to discuss what NIT adds to this existing toolset. Recent work by Timme and Lapish [28] offers an extensive review of existing IT analysis software packages. We have further complemented their work with an updated overview (**Table 1**). Of the 12 packages reviewed in Timme and Lapish [28], none satisfied simultaneously the following requirements: being applicable to both discrete and continuous data, providing means for significance testing and correction for limited sampling bias, and implementing calculation of information-theoretic measures beyond MI and transfer entropy (e.g., those based on PIDs). NIT simultaneously implements all these features.

**Table 1.** Comparison with existing information theoretic toolboxes. If the toolbox computes quantities that are defined as simple linear combinations of entropies or mutual information, for brevity we list them under entropy or mutual information.

| Toolbox | Information Measures | Data Types | Significance Testing | Probability Estimation Methods | Bias Correction | Dimensionality Reduction Methods | Language |
| --- | --- | --- | --- | --- | --- | --- | --- |
| NIT – this paper | Entropy, mutual information, transfer entropy, information breakdown, partial information decomposition, intersection information, feature information transfer | Discrete and continuous | Non-parametric | Binning (several methods)<br>Gaussian fit<br>Parametric copula (Gaussian, Clayton, student)<br>Nonparametric copula | Yes | Yes | MATLAB front-end, interfaceable with Python through MATLAB Engine API for Python |
| Information Breakdown Toolbox [25] | Entropy, mutual information, transfer entropy, information breakdown | Discrete and continuous | Non-parametric | Binning (several methods)<br>Gaussian fit | Yes | No | MATLAB |
| Gaussian Copula Mutual Information [30] | Entropy, Mutual Information | Discrete and Continuous | No | Gaussian copula | Yes | No | MATLAB and Python |
| Neuroscience Information Theory Toolbox [28] | Entropy, mutual information, transfer entropy, partial information decomposition, information transmission | Discrete and continuous | No | Binning (several methods) | No | No | MATLAB |
| JIDT [31] | Entropy, mutual information, transfer entropy | Discrete and continuous | Non-parametric | Binning<br>Kernel-based<br>Gaussian fit | Yes | No | JAVA (with Python and MATLAB wrappers) |
| FRITES [32] | Entropy, Mutual Information, transfer entropy | Discrete and continuous | Non-parametric<br>Group stats | Binning (equi-spaced)<br>Gaussian copula | Yes | No | Python |
| Inform [156] | Entropy, mutual information, transfer entropy | Discrete | No | Binning (several methods) | No | No | C (with Python, Julia, R and Mathematica wrappers) |
| Transfer Entropy Toolbox [89] | Transfer entropy | Spike trains | No | No | No | No | MATLAB |
| Trentool [152] | Transfer entropy | Continuous | Non-parametric<br>Group stats | Kernel-based | Yes | No | MATLAB |
| MuTE [157] | Transfer entropy | Continuous | Non-parametric | Binning (equi-spaced)<br>Gaussian fit<br>Kernel-based | Yes | No | MATLAB |
| ToolConnect [158] | Entropy, transfer entropy | Spike trains | No | No | No | No | C# |
| STAToolkit [159] | Entropy, mutual information | Spike trains | Non-parametric | Binning | Yes | No | MATLAB (with .mex files) |
| PyEntropy [29] | Entropy, mutual information, maximum entropy models | Discrete and Continuous | No | Binning (several methods)<br>Shrink Estimator | Yes | No | Python |
| ITE Toolbox [160] | Entropy, mutual information | Discrete and Continuous | No | Kernel-based | No | No | MATLAB and Python |
| Dit [161] | Entropy, mutual information, partial information decomposition | Discrete | No | No | No | No | Python |
| Climer and Dombeck [162] | SMGM information [163] | Discrete and Continuous | No | No | No | No | MATLAB |

The NITT Neuroscience Information Theory Toolbox [28] is, among those previously available, one of the most complete in terms of information quantities offered. However, like others listed in Table 1, it lacks limited sampling bias correction. This is not a problem when considering quantities that do not require the computation of stimulus specific distributions of neural responses, such as entropy and TE. Lack of bias corrections instead becomes a major problem for studies of coding of sensory or choice variables, as they require estimation of stimulus-related information variables that are based on calculations of stimulus-specific response probabilities. In such cases, stimulus-specific information values are dominated by the bias, if not bias corrected. A lack of bias corrections makes it impossible to meaningfully compare the amount of information carried by neural representations with different dimensionality such as spike times vs spike rates [17, 151] or single neurons vs population responses. The JIDT toolbox [31] also offers extensive sets of IT measures, although (like the NITT) it lacks methods for dimensionality reduction that are useful e.g. to apply IT to large populations. Other toolboxes [89, 152] are specialized on transfer entropy and are thus suitable for study information communication but not information encoding. Finally, some other toolboxes [30] are effective for specific distributions of neural activity, such as the case of Gaussian interactions which are relevant for mass measures of activity, but are difficult to apply to measures with single cell resolution for which statistics and interactions are not well described by Gaussian distributions.

We made an effort to improve computational performance in NIT, designing it to maximize efficiency and scalability. This optimized design strategy resulted in fast computational times compared with other state-of-the-art open access codes. We benchmarked our toolbox against NITT [28] on a single MI calculation, with bootstrap null distribution estimation, obtaining on average 50 times faster computation times with NIT compared to NITT.

NIT does have limitations, which we plan to address in ongoing and future updates. NIT still lacks computation of useful quantities, such as maximum entropy (ME) models, which are useful to determine the order of interactions among neurons [35, 153]. ME models are present in some specialized toolboxes [29]. Further, NIT includes standard and widely used non-parametric hypothesis testing methods, but does not yet include group statistics, which to the best of our knowledge among information theoretic toolboxes has been only implemented in FRITES [32]. However, the output of NIT analyses can be easily used as input to group statistics toolboxes [32]. Further, the study of PID is a burgeoning field with many measures and advances being elaborated [49, 80, 154, 155]. While NIT implements some of the most established PID quantities, it will be important to keep it updated to include more PID developments and to interface with new PID software.

### Validations and recommendations for the analysis of calcium imaging

The methods presented in NIT are applicable to any kind of neuroscience recordings, both discrete and continuous. Given that the plug-in binning estimators presented here have been extensively and successfully validated on electrophysiological data (from spike trains, to LFP and EEG), in this study we focused on validating the information-theoretical analysis of 2P calcium imaging data. 2P imaging signals are potentially more challenging than electrophysiological ones to analyze with information theory, because of the lower SNR and temporal resolution. Moreover, the problem of how to recover from calcium traces as much information as possible about external stimuli or about the underlying spiking activity of the imaged neurons has not been systematically studied yet.

We addressed these issues using a thorough analysis of synthetic calcium imaging traces, generated through a biophysically plausible single-compartment model of cytosolic calcium dynamics. Specifically, we assessed the effect of the calcium indicator (GCaMP6f vs GCaMP6s), imaging frame rate, SNR, response profile shape, and spike rate modulation by the stimulus on the stimulus information computed from the simulated calcium signal. We found that estimates of MI from the ΔF/F signal depended relatively weakly on the imaging frame rate and SNR. However, the amount of MI that could be obtained from calcium fluorescence traces is the temporal shape of the neuronal response. A tonic neuronal response transfers more information in the calcium signal compared to a phasic one, particularly when using an indicator with slow decay time and high dynamic range (GCaMP6s). We have further observed that, when the neuron encodes the stimulus in a phasic way at high firing rates, the calcium signal can occasionally encode more stimulus information than the time-averaged spike rate (**Figure 4**). The reason for this counterintuitive finding is that in this condition spiking activity is concentrated within a limited time interval and thus knowledge of when spike times are more informative adds information, and that the nonlinearities of calcium dynamics emphasize the signal in this high-firing high-information region and deemphasize the signal in the low-firing low information region, thereby achieving more information than the time-average spike rate which instead weighs all spikes equally regardless of when they were fired.

Furthermore, we have proposed a new single-trial calcium metric, based on the inversion of the forward model that we have used for the generation of synthetic calcium traces, for the estimation of calcium concentration in the cell given a ΔF/F trace. This approach was inspired previous work [164] inferring action potentials by building an inverse model of membrane potential from calcium imaging signals. We assessed the performance of this single-trial calcium metric for computing information from calcium data, and we compared it with other widely used strategies for quantification of single trial ΔF/F responses. We found that, across all simulation conditions examined, the newly proposed estimated calcium and the linear deconvolution of the ΔF/F trace with a decaying exponential were the two single trial calcium response quantification that allowed to extract more information (about external stimuli or about the underlying spike rates). Other considered quantifications of single trial calcium responses (max ΔF/F, mean/integral ΔF/F, OASIS) extracted less information. These results were confirmed on experimental data coming from four independent datasets – including both GCaMP6f and GCaMP6s signals simultaneously acquired on individual cells together with juxtasomal electrophysiological recordings. Careful choice of single-trial quantifications of calcium signals can, thus, significantly increase the amount of information retrieved, and we propose a new and efficient metric to do so.

Importantly, we compared different information computation methods, all implemented in NIT, to compute information from calcium data. We found that the non-parametric copula-based estimator for mutual information [71] was the one working best, outperforming both binned estimators and parametric Gaussian copulas in terms of data robustness and accuracy of the estimation. While the non-parametric copula comes at the expense of major increase of computing time, it should be recommended for calcium data whenever its computation is practically feasible.

A result of importance of our simulations and real data analysis was that, when proper quantification and algorithms were applied, we could recover surprisingly large amounts of information from calcium imaging. In simulations, the amount of stimulus information obtained from realistically simulated calcium imaging traces was > 50% of the stimulus information encoded in the simulated spike trains when effective single-trial calcium metric were applied (Fig 5B). In both simulated and real data, a relatively large amount of information about the underlying spike rate could be recovered from the calcium traces when using appropriate calcium metrics and algorithms (Fig 5D,7). These results illustrate the power of calcium imaging for studying population activity and the importance of coupling it with advanced information theoretic and signal extraction methods.

Climer and Dombeck [162] have recently discussed the application to calcium imaging of a specific information metric termed SMGM. This metric has been first introduced by Skaggs et al. [163] for electrophysiological data and is often used in the literature for hippocampal place field quantification. It has been shown [165] that, when applied to spike trains, the SMGM metric approximates well the full information content of a spike train only when the average number of spikes per trials is much smaller than 1 (i.e. very low firing rates or very short time windows) and that the correlations between spikes are small enough so that the firing statistics is close to that of a Poisson process. Using the SMGM metric with the ΔF/F signal as a proxy of the information carried by the underlying spike rates rate additionally assumes that a constant proportionality exists between the firing rate and fluorescence signal for a given indicator. However, there are known non-linearities between spike rate and fluorescence. Using MI to extract information from calcium traces as a proxy of information from spike rates does not require the assumption of a linearity between spike rates and calcium fluorescence, because MI is insensitive to monotonic non-linearities in the transformation between variables, and it does not require the assumption that neuron fire at very low rates with Poisson statistics. Based on these considerations, we recommend application of SMGM to estimate information from calcium imaging data only when there is an expectation of linearity between spike rates and calcium responses and of very low firing rates of neurons. Estimations made using MI are instead valid and applicable under more general circumstances.

### Conclusions

Overall, our toolbox provides a comprehensive set of information theoretic measures applicable to any kind of neuroscience data.

## Materials and methods

### Details of the performed parametric simulation sweep

Below are listed the values considered for each of the variables considered in the parametric sweep of simulations of neural activity and calcium imaging traces.

- Imaging frame rate: 5, 10, 100 Hz.
- SNR: 5, 9, 15.
- PSTH shape: Tonic (gaussian-shaped with peak at 0.25s over a 1s trial duration, standart deviation 0.01 s), phasic (uniform distribution over time).
- Stimulus modulation of neuron mean firing rate:

- [1 Hz – 2 Hz]: *Low MI, Low SR*
- [12 Hz – 13 Hz]: *Low MI, High SR*
- [2 Hz – 12 Hz]: *High MI*
- Indicator: GCaMP6f, GCaMP6s.
- Number of trials per stimulus: [5,10,20,30,40,50,60,80,100,200,400].

### Mutual Information (Direct plug-in method)

*MI*(*S;R*) has been calculated using Equation (1), where the marginal and joint probabilities have been calculated by simply counting the number of occurrences of the discrete values of *R* and *S* across repeated presentations of the stimulus. If variables *R* and *S* were continuous, they were discretized using binning routines. The binning strategy and number of bins used for each specific analysis using direct plug-in method are reported in the main text, together with the use of bias correction method used for the specific analysis.

### Mutual Information (Non-Parametric Copula)

We estimated the mutual information between two variables *R* and *S* using the nonparametric copula approached presented in [71]. Copula is defined as the probability function between the CDF’s of the marginal variables *U_R_*~CDF(*R*) and *U_S_*~CDF(*S*) and it captures the general correlation structure of the joint density function between variables. To compute the mutual information I(*R*; *S*), we use the fact that it is related to the copula entropy as:

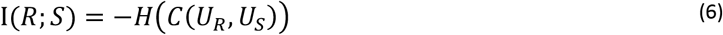

Where *C*(*U_R_, U_S_*) is the joint density function of CDF variables *U_R_* and *U_S_*. To compute the copula density, we used the same analytic solution for a local likelihood kernel estimation of the CDF values after optimizing the bandwidth using a genetic optimization developed in Safaai et al. [71]

We then estimated the copula density over the whole space of CDF’s (*U_R_, U_S_*) using the optimized kernels and on a grid of size k which defines the resolution of density estimation. We normally used k=50 or k=100 in our calculation and the change didn’t make significant difference on our results.

After estimating the copula density on the grid, we generated correlated samples of data by first computing the conditional cumulative copula density by integrating the copula density over the grid:

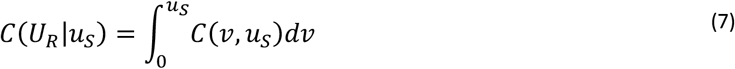

Which is a uniform distribution. Using the fact that the marginal distribution of a CDF distribution is uniform, the 2-dimensional correlated samples can be generated as follows:

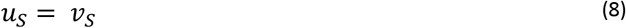

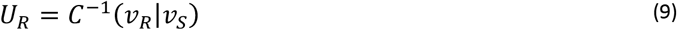

Where *v_R_* and *v_S_* are independent samples from the uniform distribution 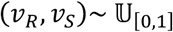. We then used these samples to estimate the copula entropy, using classical Monte-Carlo approach after expressing the entropy as the expectation over copula density 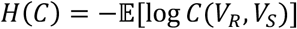. For the case in which one of the variables is discrete, we first transform the variable into the continuous domain by adding an appropriate noise which as it was shown in Safaai et al. [71].

### Mutual Information (Parametric Copula)

We implemented several algorithms for mutual information estimation using parametric copulas that have been introduced in neuroscience [30, 70]. Full details are contained in the software documentation. In brief, we adapted our algorithms from those of Ref (Onken and Panzeri, 2016). For continuous margins, we provide implementations of the normal and the gamma distributions. For discrete margins, we provide the Poisson, binomial and negative binomial distributions. We provide the Gaussian, student and Clayton bivariate copula families as well as rotation transformed Clayton families.

### Generation of synthetic calcium imaging traces

#### Convolution with a double exponential kernel

Fluorescent signal was generated as a convolution of the input spike train with a double exponential kernel in the form:

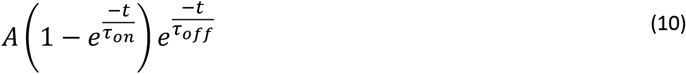

Chen et al.[122] report values of peak amplitude, peak time and half decay time for both GCaMP6f and GCaMP6s in mouse V1 in vivo experiments. Those values are related to the constants *A, τ_on_* and *τ_off_* defined above, and have been defined through an iterative optimization to generate a double exponential kernel with the same peak amplitude, peak time and half decay time than reported in literature. Values used of the three constants for the two indicators are reported in **Supplementary table 1**.

**Supplementary table 1.** Used constants for the synthetic trace generation through a double exponential kernel

|  | GCaMP6f | GCaMP6s |
| --- | --- | --- |
| A [-] | 0.39 | 0.51 |
| $\tau_{on}$ [s] | 0.03 | 0.14 |
| $\tau_{off}$ [s] | 0.09 | 0.37 |

Gaussian white noise with given standard deviation is added to the convolved trace to generate the synthetic calcium imaging trace with given SNR.

#### Biophysically plausible SCM

Evolution of cytosolic calcium concentration [Ca] is modelled trough the following differential equation [120]:

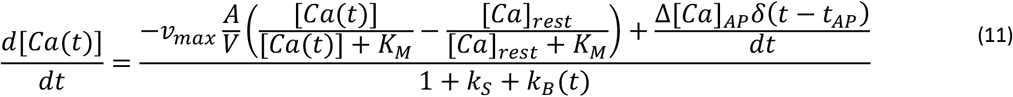

Where *v_max_* is the maximum efflux rate per unit area of the cell membrane, *A* is the membrane area, *V* is the compartment volume, *K_M_* is the concentration at which extrusion is half maximal, Δ[*Ca*]_*AP*_ is the amount of calcium intake following an action potential, *δ* is Dirac’s delta, *t_AP_* are the times of action potentials, *k_s_* is the binding ratio of the endogenous [Ca] buffers, and *k_B_* is the binding ratio of the exogenous buffers (the indicator itself). The latter is not a model constant and, for the case of cooperative binding, is defined as [166]:

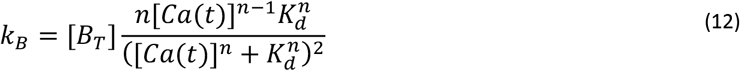

Where [*B_T_*] is the concentration of the indicator, *n* is the Hill coefficient, *K_d_* is the dissociation constant of the indicator.

Equation (11) contains two non-linear terms: a saturable mechanism for calcium extrusion from the cytoplasm (first term at the numerator on the right-hand side). Measured values of *v_max_* are hardly available in literature. It is more common to find estimates of the extrusion rate *γ* in case of a linear extrusion mechanism (*Ca*(*t*)_*out*_ = *γ*([*Ca*(*t*)] – [*Ca*]_*rest*_))[120]. We have thus specified *v_max_* so that the extrusion rate would match *γ* = 1200 [1/*s*] in the surroundings of [*Ca*(*t*)] = [*Ca*]_*rest*_.

Time integration of Equation (11) allows to obtain the time trace of free cytosolic calcium in the cell. The concentration of indicator bound calcium [*CaB*(*t*)] has been obtained through integration of:

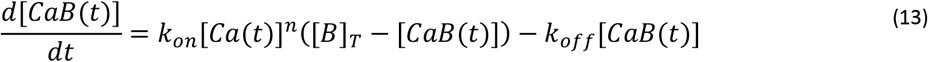

Where *k_on_* and *k_off_* are the association/dissociation rates.

Once known the fraction of calcium-bound indicator, fluorescence is generated through a linear model [166]:

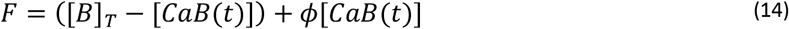

Where the constant Φ is indicator specific and has been tuned to experimental data.

The value of baseline fluorescence *F*_0_, in resting state, steady conditions, is calculated from the resting state indicator-bound concentration using Equation (14)[166]:

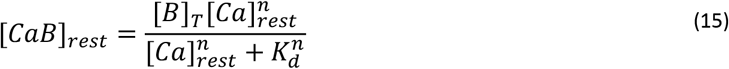

The model then returns the normalized fluorescence, with the addition of white noise term:

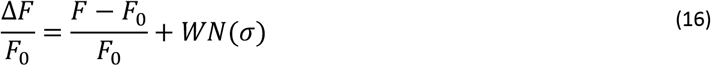

The standard deviation of the white noise has been specified to match the desired SNR for a given synthetic trace.

**Supplementary table 2.**
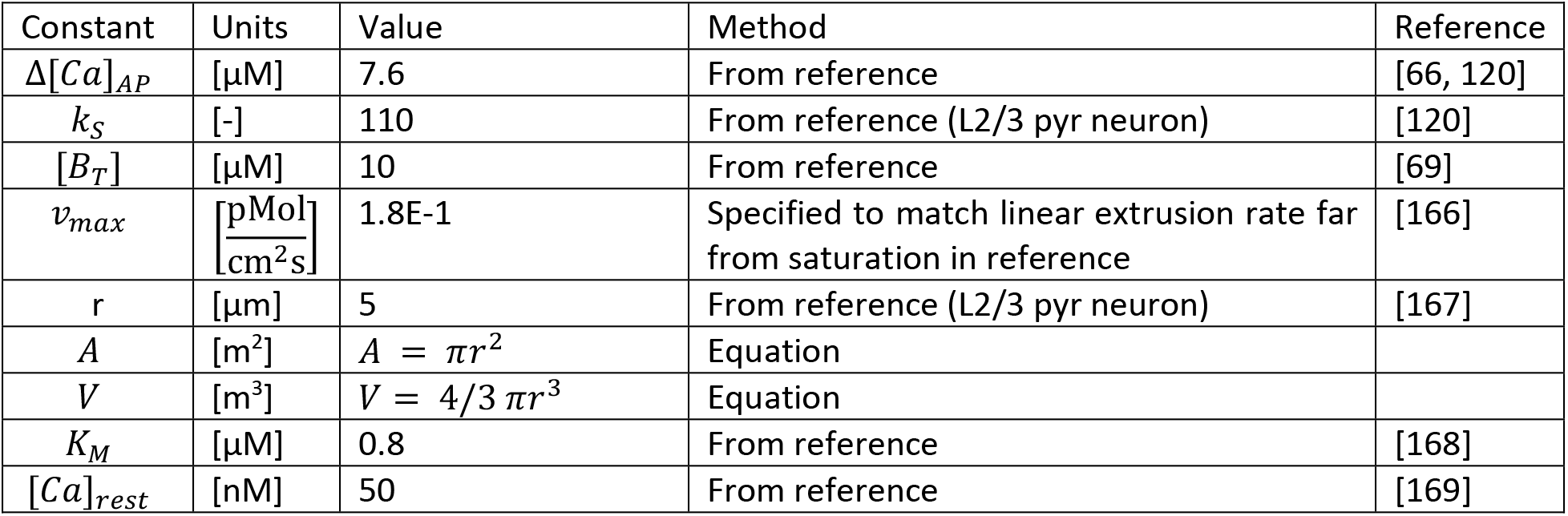
Model constants used in the SCM. These constants were independent on the indicator.

| Constant | Units | Value | Method | Reference |
| --- | --- | --- | --- | --- |
| $\Delta[Ca]_{AP}$ | [μM] | 7.6 | From reference | [66, 120] |
| $k_s$ | [-] | 110 | From reference (L2/3 pyr neuron) | [120] |
| $[B_T]$ | [μM] | 10 | From reference | [69] |
| $v_{max}$ | $\frac{[pMol]}{[cm^2s]}$ | 1.8E-1 | Specified to match linear extrusion rate far from saturation in reference | [166] |
| $r$ | [μm] | 5 | From reference (L2/3 pyr neuron) | [167] |
| $A$ | [$m^2$] | $A = \pi r^2$ | Equation | |
| $V$ | [$m^3$] | $V = 4/3 \pi r^3$ | Equation | |
| $K_M$ | [μM] | 0.8 | From reference | [168] |
| $[Ca]_{rest}$ | [nM] | 50 | From reference | [169] |

**Supplementary table 3.** Indicator specific constant used in the SCM. These constants were indicator-specific and have been determined through fitting the model on experimental data.

| Constant | Units | Value GCaMP6f | Value GCaMP6s | Method | Reference |
| --- | --- | --- | --- | --- | --- |
| $n$ | [-] | 2.47 | 2.93 | Fit to experimental data | |
| $K_d$ | [nM] | 375 | 144 | From reference | [122] |
| $k_{on}$ | [Hz/M<sup>n</sup>] | $k_{on} = k_{off}/K_d^n$ | | From reference | [166] |
| $k_{off}$ | [Hz] | 5.16 | 0.5 | Fit to experimental data | |
| $\Phi$ | [-] | 15.01 | 62.72 | Fit to experimental data | |

#### Fitting of the SCM to experimental data

Fitting of the single-compartment model is done in the following way (separately for each indicator considered). Among the three variables that are fit to data (Φ, *n, k_off_*), the first one is optimized first – and independently from the other two – so that saturated indicator reaches the dynamic range reported in [122]. This is possible due to the fact that *n* and *k_off_* do not impact the steady state brightness of the indicator, but only its dynamics. Simultaneous 2-photon imaging and cell-attached electrophysiology data[170] are then used to define the kinetics of the indicator binding/unbinding and its cooperativity. Given the experimentally measured spike train, and SNR of the experimental fluorescent trace, we have optimized *n* and *k_off_* to reduce the root square error between the generated synthetic calcium trace and experimental data. Dataset ‘data_20120521_cell5_007.mat’ has been used for GCaMP6f tuning, while ‘data_20120515_cell1_006.mat’ has been used for GCaMP6s.

### Definition of spiking activity metrics based on ΔF/F

#### Max ΔF/F

Values of peak ΔF/F over a defined post stimulus time interval have been calculated as follows:

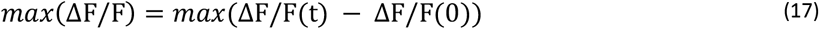

#### Mean/integral ΔF/F

Values of mean ΔF/F over a defined post stimulus time interval have been calculated as follows:

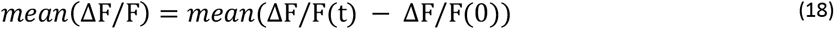

It should be noted that, throughout the text, we refer to this metric as Mean/integral ΔF/F. The reason for this is that the mean and integral are related by a constant linear scaling and are de facto equivalent in information-theoretical terms. The full dataset attached to this paper contains also separate analysis for *integral*(ΔF/F), showing identical performance to *mean* (ΔF/F’).

#### Estimated Calcium

This metric of spiking activity based on the two photon imaging recordings is based on the inversion of the forward model detailed in section Biophysically plausible SCM. The inversion, calculating thus [Ca] from the ΔF/F assumes that the binding/unbinding happens at chemical equilibrium. In this condition, for cooperative binding, we can write the relation between [CaB] and [Ca] as:

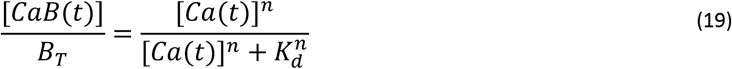

Deriving both left and right-hand side:

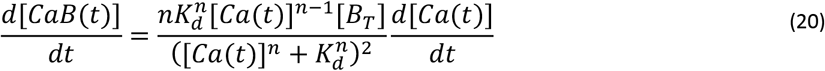

Assuming that the generated fluorescence is a linear combination of the fractions of calcium-free [B] and calcium-bound [CaB] indicator we can write the following:

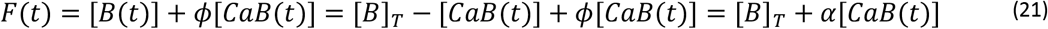

Where *α* = *ϕ* – 1. Baseline state fluorescence, thus, is:

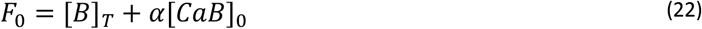

Combining Equations (21) and (22) we have that:

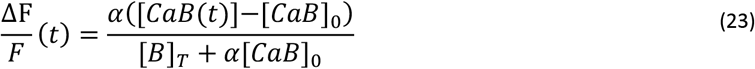

Deriving both left and right-hand side of Equation (26) with respect to time:

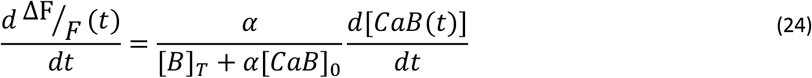

Combining Equations (24) and (20):

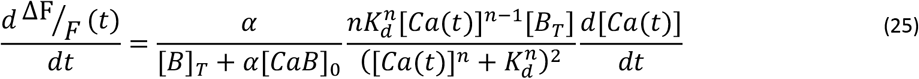

Given the time trace of fluorescence, equation (28) can be used to solve for [*Ca*(*t*)] once [Ca(0)] and [*CaB*]_0_ are known. These values have been defined through the following educated guesses. The baseline calcium-bound indicator concentration [*CaB*]_0_ is taken as the steady state equilibrium concentration when [*Ca*] = [*Ca*]_*rest*_ (using Equation (19)).

In order to estimate the initial concentration of free calcium in the cell we used the following approach. Combining equations (23) and (19) we obtain:

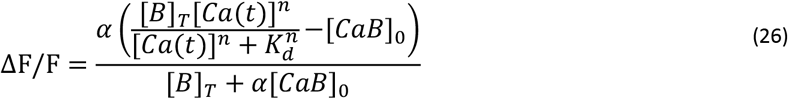

Given ΔF/F at time zero, iterative solution of equation (26) for [Ca(0)]. This sets the initial conditions for time integration of equation (28).

The obtained time trace is finally deconvolved through a single decaying exponential kernel with time constant equal to the reciprocal of the unbinding rate of the indicator koff. The mean of the deconvolved trace is reported as the estimated [Ca].

#### Linear deconvolution

The ΔF/F trace has been deconvolved with a decaying exponential with a decaying time constant *τ_off_*. The reported value of the deconvolved signal over the post stimulus time interval has been calculated as:

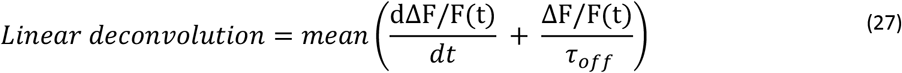

Where the values of *τ_off_* (*τ_off_* = 2*s* for GCaMP6s and *τ_off_* = 0.5*s* for GCaMP6f) have been estimated from the decay of time traces for GCaMP6 indicator reported in Chen et al.[122]

#### OASIS

Time trace of ΔF/F has been deconvolved using MATLAB implementation of OASIS [137] (https://github.com/zhoupc/OASIS_matlab). We have used the second order auto-regressive thresholded implementation of the algorithm. This implementation imposes a minimum threshold for the deconvolved trace, effectively filtering out spurious deconvolved activity. The parameters of the auto-regressive model, the value of the threshold, as well as the SNR levels were estimated by internal functions of the toolbox. The returned value of OASIS metric over a post-stimulus window has been calculated as:

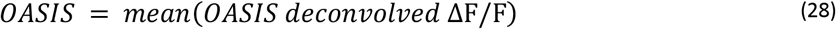

In order to avoid potential issues in using OASIS to deconvolve traces of limited duration, the time traces of ΔF/F extended for a total duration of 10s, of which the first second was stimulus modulated and the remaining part had a constant baseline SR of 0.5 Hz.

### Definition of preparatory activity in motor cortex

Stimulus and choice instantaneous information were computed using mutual information between those variables and the neural activity over time, resulting in values of information over the trial duration. Mutual information and intersection information were computed using the direct plug-in method for computational tractability of such a large dataset. Neural activity was binned in 2 equally populated bins for every timestep.

## Software availability

NIT source code, documentation, installation instructions and tutorials can be downloaded from the following repository: https://gitlab.com/rmaffulli/nit. Software for the realistic calcium imaging simulations can be downloaded from the following repository: https://gitlab.com/rmaffulli/casim

## Data availability

Simultaneous calcium imaging and electrophysiological recordings used for Figure 7 taken from Refs [119, 122, 141] can be obtained as specified in these publications. Simultaneous calcium imaging and electrophysiological recordings used for Figure 7 taken from Ref [142] can be obtained from the corresponding authors upon reasonable request.

## Author contributions

RM: conceptualization, data curation, formal analysis, funding acquisition, investigation, methodology, software, validation, visualization, writing – original draft preparation, review & editing. MAC: conceptualization, data curation, formal analysis, investigation, software, visualization, writing – original draft preparation, review & editing. MC: software, writing – review & editing. SZ: investigation, data curation. TF: supervision, funding acquisition, writing – review & editing. HS: supervision, software, writing – review & editing. SP: conceptualization, supervision, funding acquisition, writing – review & editing.

## Acknowledgements

The authors would like to thank Dr Mariangela Panniello for helpful feedback and discussions.

## Supplementary material

### Supplementary tables

**Supplementary table 4.** Compatibility matrix between information-theoretic quantities in NIT and applicable bias correction strategies.

| Information quantity | Allowed bias correction methods |
| --- | --- |
| Mutual Information (direct method) | Quadratic Extrapolation, Panzeri-Treves, Bootstrap correction, BUB |
| Mutual Information (non-parametric copula) | Bootstrap correction |
| Mutual Information Breakdown | Quadratic Extrapolation, Panzeri-Treves, Bootstrap correction |
| Transfer Entropy | Quadratic Extrapolation, Panzeri-Treves, Bootstrap correction |
| Partial Information Decomposition | Linear Extrapolation, Quadratic Extrapolation, Bootstrap correction |
| Intersection Information | Linear Extrapolation, Quadratic Extrapolation, Bootstrap correction |
| Feature Information Transfer | Linear Extrapolation, Quadratic Extrapolation, Bootstrap correction |

**Supplementary table 5.** Table of p-values and effect sizes ω^2^ for data in **Figure 3C.** Data have been analyzed using a separate three-ways ANOVA (considering PSTH shape, FR and SR as grouping variables) for each information level.

|  | PSTH shape | FR | SNR |
| --- | --- | --- | --- |
| Low MI, low SR | p-val = 2.24e-16<br>$\omega^2 = 7.00e-02$ | p-val = 3.97e-04<br>$\omega^2 = 1.40e-02$ | p-val = 7.21e-02<br>$\omega^2 = 3.32e-03$ |
| Low MI, high SR | p-val = 8.11e-48<br>$\omega^2 = 2.08e-01$ | p-val = 4.05e-01<br>$\omega^2 = -1.70e-04$ | p-val = 4.09e-01<br>$\omega^2 = -1.85e-04$ |
| High MI | p-val = 0<br>$\omega^2 = 8.89e-01$ | p-val = 9.30e-17<br>$\omega^2 = 7.99e-03$ | p-val = 5.00e-14<br>$\omega^2 = 6.54e-03$ |

**Supplementary table 6.**
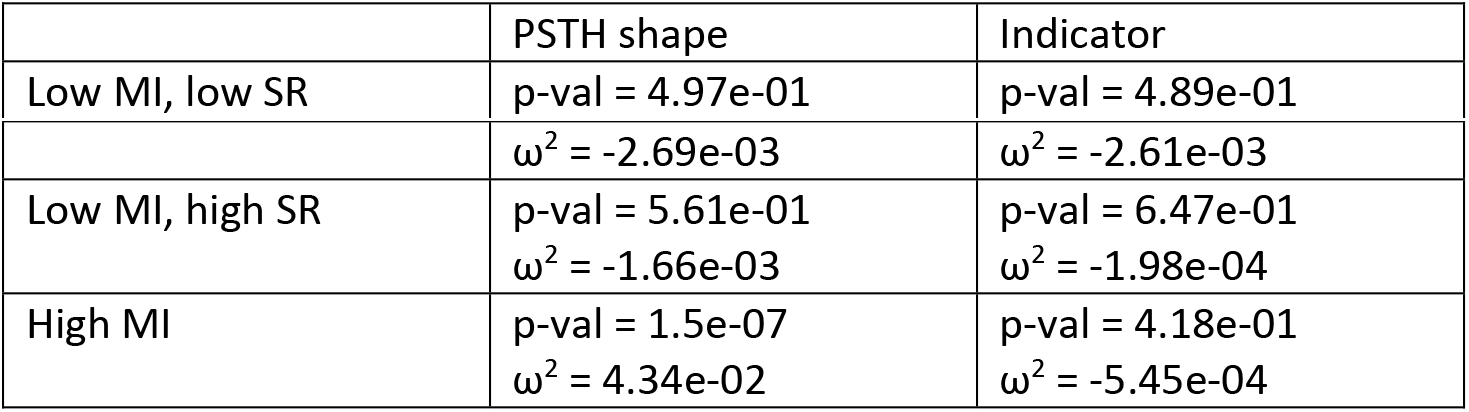
Table of p-values and effect sizes ω^2^ for data in **Figure 3D.** Data have been analyzed using a separate two-ways ANOVA (considering PSTH shape and calcium indicator as grouping variables) for each information level.

|  | PSTH shape | Indicator |
| --- | --- | --- |
| Low MI, low SR | p-val = 4.97e-01 | p-val = 4.89e-01 |
| | $\omega^2 = -2.69\text{e-}03$ | $\omega^2 = -2.61\text{e-}03$ |
| Low MI, high SR | p-val = $5.61\text{e-}01$<br>$\omega^2 = -1.66\text{e-}03$ | p-val = $6.47\text{e-}01$<br>$\omega^2 = -1.98\text{e-}04$ |
| High MI | p-val = $1.5\text{e-}07$<br>$\omega^2 = 4.34\text{e-}02$ | p-val = $4.18\text{e-}01$<br>$\omega^2 = -5.45\text{e-}04$ |

### Supplementary figures

**Supplementary figure 1.**
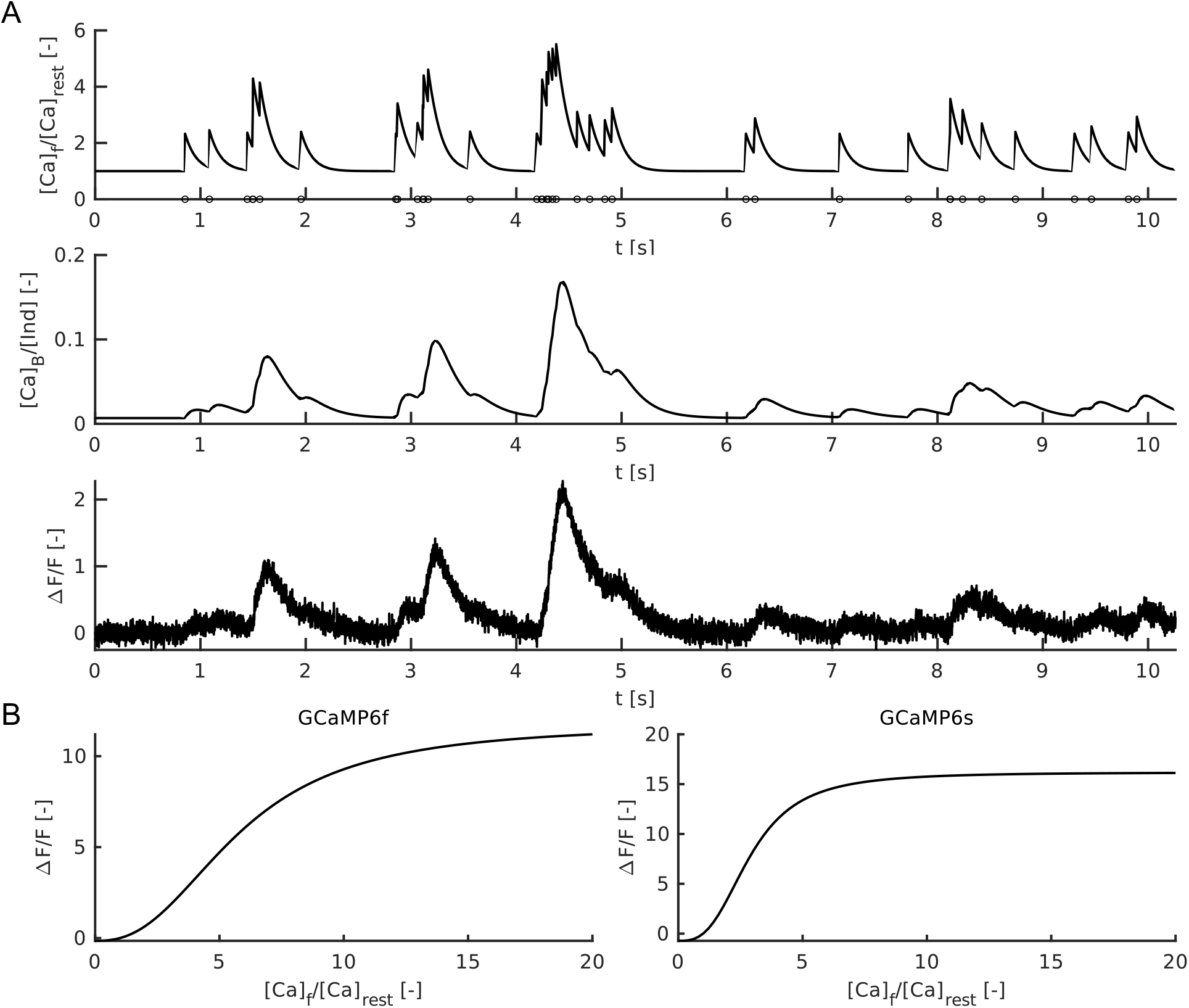
Generation of fluorescence trace in the Single Compartment Model. (**A**) Top: simulated trace of relative levels of free calcium concentration in the cytoplasm with respect to resting state levels. Circles represent action potentials. Middle: simulated trace of the fraction of GCaMP indicator bound to calcium. Bottom: fluorescent trace resulting from the fractions of calcium-bound and calcium-free indicator. (**B**) Relation between generated fluorescence and free calcium concentration in the cytoplasm in chemical equilibrium conditions for both GCaMP6f and GCaMP6s in the used model.

**Supplementary figure 2.**
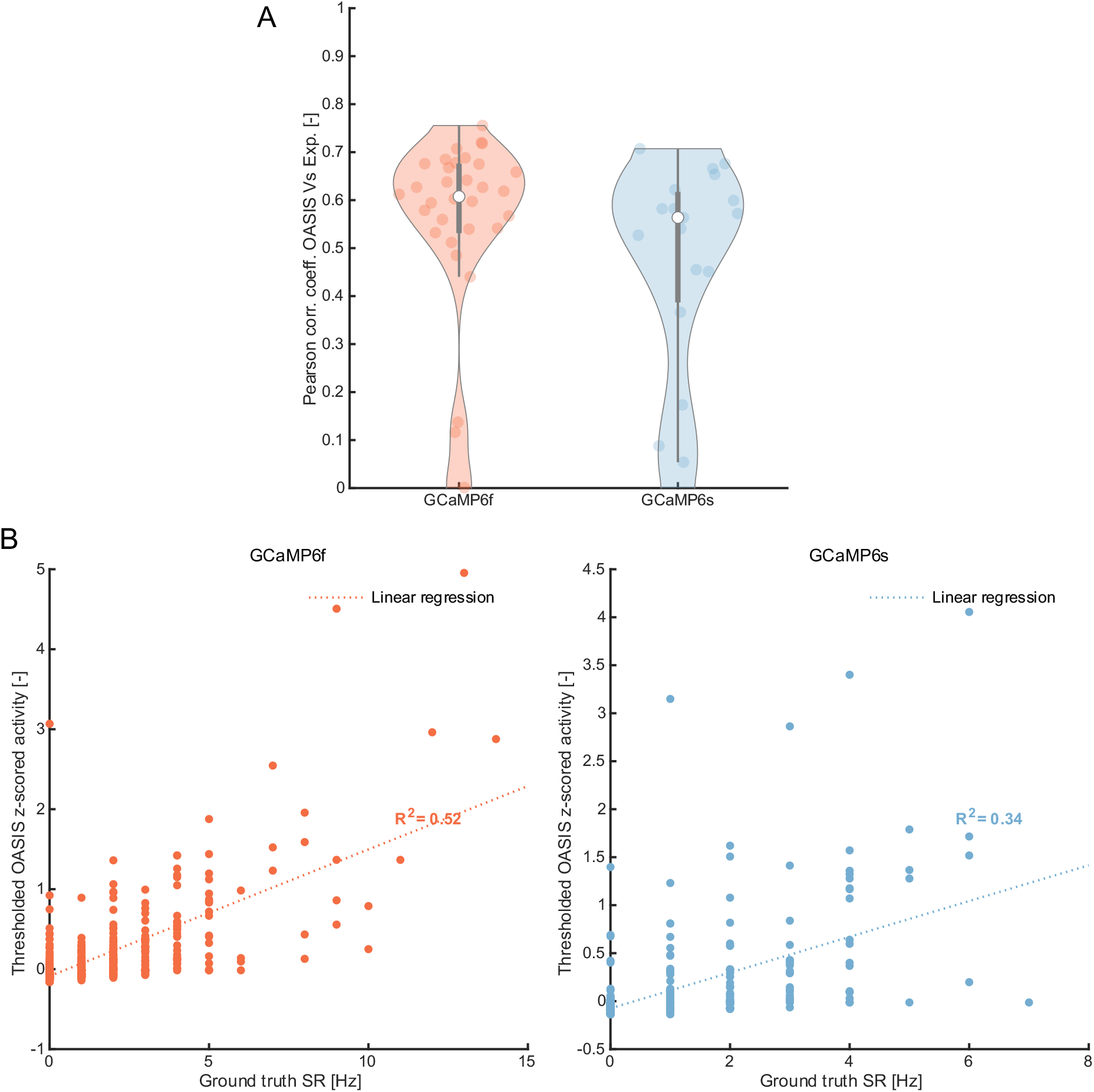
Performance of OASIS on experimental calibration dataset[170] with simultaneous calcium imaging and electrophysiology. (**A**) Pearson’s correlation coefficient between real and inferred spiking activity using 2^nd^ order auto-regressive (AR) thresholded OASIS[137] (see Materials and methods). (N = 34 for GCaMP6f, N = 19 for GCaMP6s). (**B**) Relation between z-scored inferred spiking activity in OASIS and ground truth spike rate on 1 s long windows selected randomly over the entire experimental acquisition (50 random windows per each experimental trace N = 1700 for GCaMP6f, N = 950 for GCaMP6s). Experimental data for this dataset are publicly available at: https://crcns.org/data-sets/methods/cai-1

**Supplementary figure 3.**
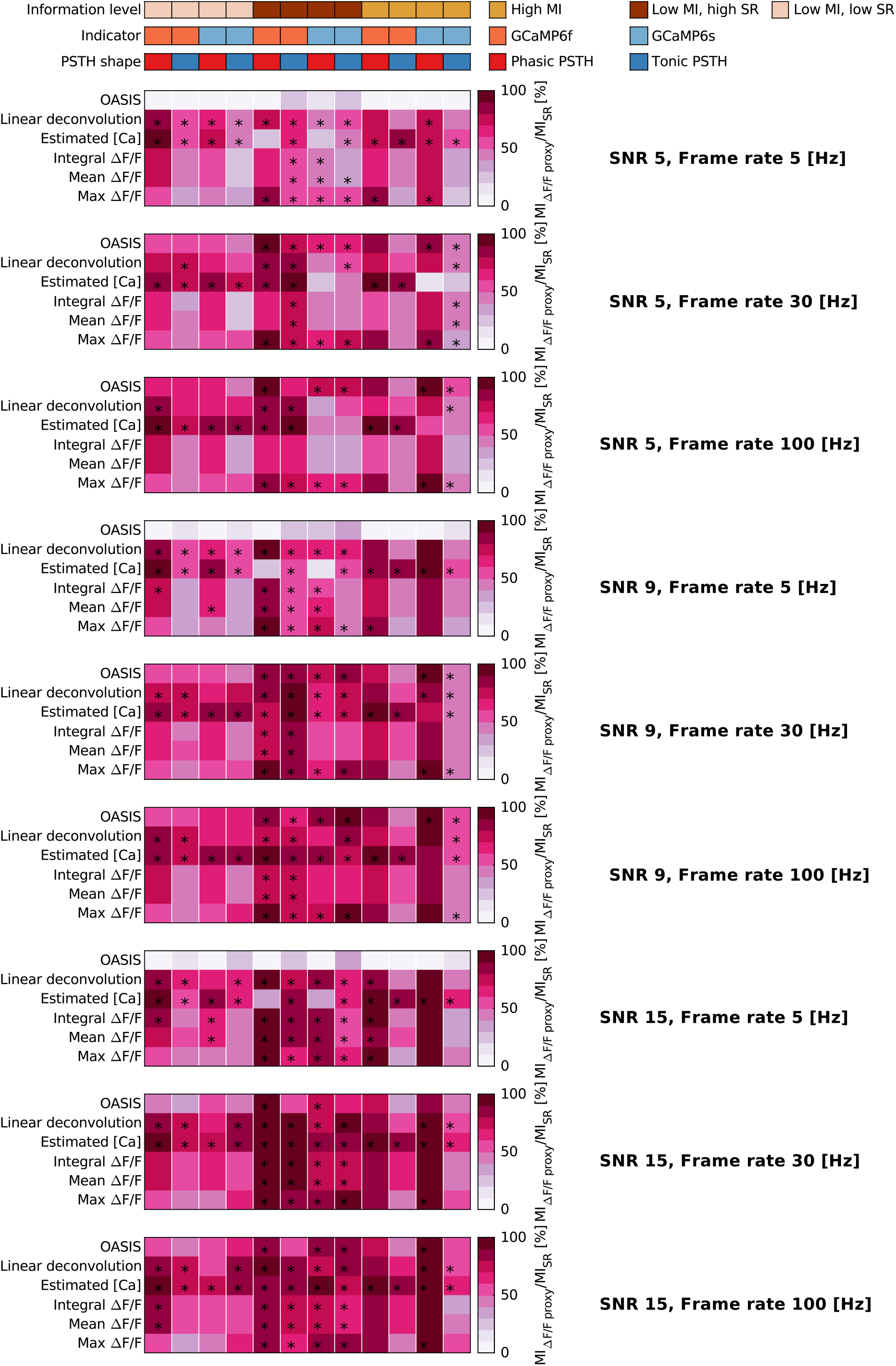
Information content in ΔF/F traces with respect to SR code. Percentage of stimulus information retrieved by each ΔF/F metric with respect to the one contained in spike rate, in all conditions of the parametric sweep considered in the study. Values represent the average over 50 simulations. For each combination of frame rate, SNR, information level, indicator and PSTH shape, the * symbol marks the metrics with non statistically different mean (p > 0.05 Bonferroni corrected Kruskal-Wallis multiple comparison test) from the best performing metric at those conditions. Best performing metric is defined as the one returning the highest mean stimulus information. All data in the figure refer to simulated traces. Mutual information is evaluated using plug-in method.

**Supplementary figure 4.**
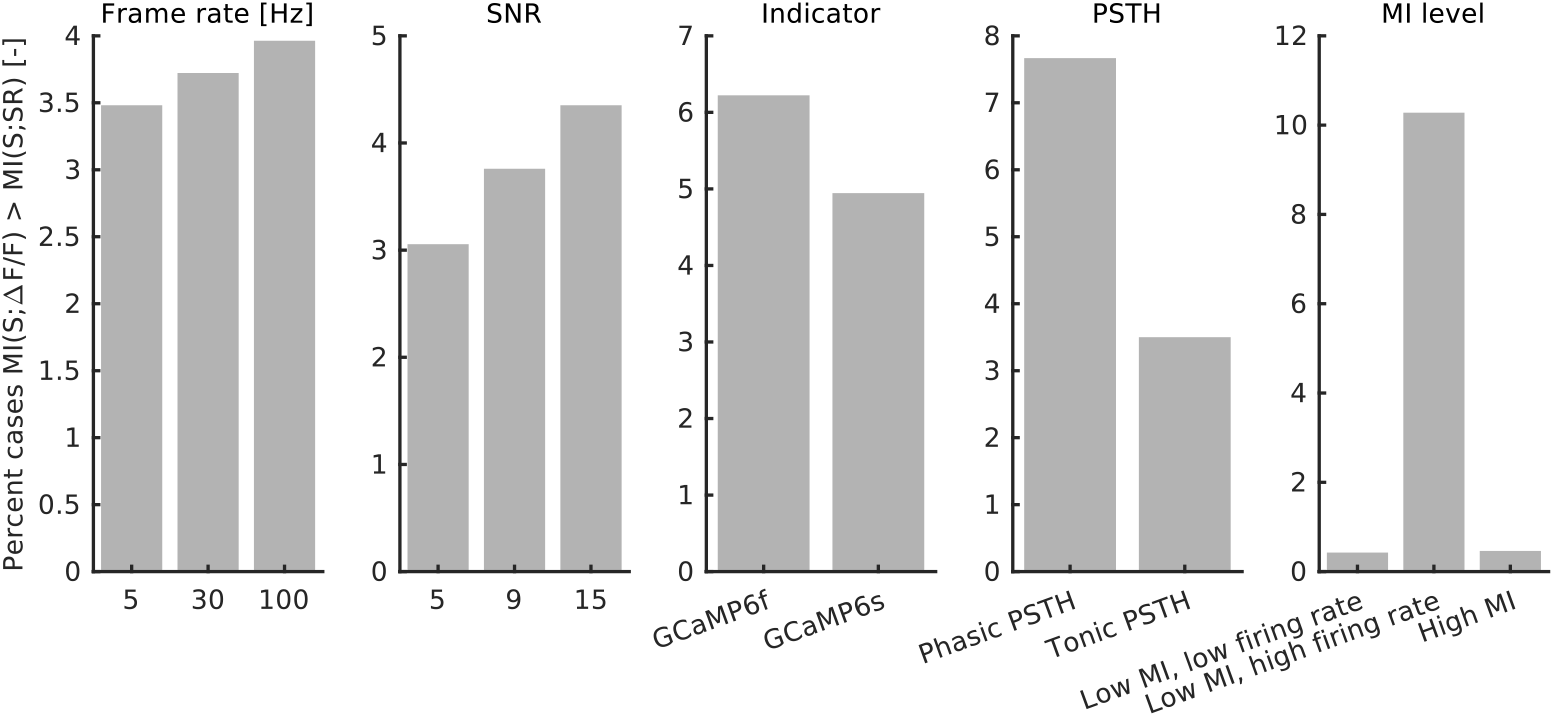
Where is MI in max ΔF/F higher than MI in SR. Percentage of cases, across all conditions investigated in the parametric sweep, where MI in max ΔF/F has been found to be higher than the stimulus information in the spike rate code.

